# Peptidergic neurons with extensive branching orchestrate the internal states and energy balance of male *Drosophila melanogaster*

**DOI:** 10.1101/2024.06.04.597277

**Authors:** Yutong Song, Tianmu Zhang, Xiaoli Zhang, Tae Hoon Ryu, Kyle Wong, Zekun Wu, Yanan Wei, Justine Schweizer, Khoi-Nguyen Ha Nguyen, Alex Kwan, Kweon Yu, Woo Jae Kim

## Abstract

Neuropeptide SIFamide (SIFa) neurons in *Drosophila melanogaster* have been characterized by their exceptionally elaborate arborization patterns, which extend from the brain into the ventral nerve cord (VNC). SIFa neurons are equipped to receive signals that integrate both internal physiological cues and external environmental stimuli. These signals enable the neurons to regulate energy balance, sleep patterns, metabolic status, and circadian timing. These peptidergic neurons are instrumental in orchestrating the animal’s internal states and refining its behavioral responses, yet the precise molecular underpinnings of this process remain elusive. Here we demonstrate that SIFa neurons coordinate a range of behavioral responses by selectively integrating inputs and outputs in a context-dependent manner. These neurons engage in a feedback loop with sNPF neurons in the ventral nerve cord, modifying behaviors such as long mating duration (LMD) and shorter mating duration (SMD). Furthermore, SIFa neurons receive essential inputs from neuropeptides Dsk, sNPF, and dilp2, which regulate interval timing behaviors. Activating SIFa neurons leads to reduced mating duration and increased food intake, while deactivating them reduces food intake. Overall, these findings demonstrate the importance of SIFa neurons in absorbing inputs and turning them into behavioral outputs, shedding light on animal’s intricate behavioral orchestration.

## INTRODUCTION

The brain’s activity is always changing, jumping from one state to another. This has a big impact on not only our neural responses to sensory inputs but also on our ability to process this huge amount of information, make decisions, and behave in the proper way^1,2^. Our decisions are frequently accompanied by ‘internal states (or ρε states)’ like motivation, arousal, desire, or emotion. Internal states have common characteristics, including persistence and scalability^3^. However, very little is known about how such internal states are recorded in animal brains and whether they impact behavioral decision-making directly or indirectly.

Internal states in reproductive social behaviors, such as mating and fighting, have been intensively studied^3–6^. Genetic methods in flies have enabled for the discovery of discrete neuronal groups expressing dopamine^7^, 5-HT^8^, or octopamine^9^, all of which control aggressive behavior. The male courtship circuitry of the fruit fly is one of the most intensively researched neural systems for understanding how the brain governs an intrinsic reproductive social behavior^5,6,10^. Nutrient homeostasis also creates unique internal states of the brain, which influence an animal’s food preference^11,12^. The shift between internal states and their influence on behavioral decision-making have been studied in courtship, aggressiveness, and nutrient-specific appetites, but how these shifts are encoded and combined to modify specific choice is unclear^3,11–17^.

Interval timing is a specific timescale that is measured precisely within the range of seconds to minutes^18^. Humans and other animals use interval timing for foraging, multi-step arithmetic, and decision-making^18–24^. Recent research suggests the brain encodes time as temporal changes in network states^25^. Observations into the interplay between sensory inputs and the impact of internal states like attention have been uncovered by psychophysical measures of interval timing^26^.

The mating duration of male fruit flies is a suitable model for studying interval timing and it could change based on internal states and environmental context. Previous studies by our group^27–30^ and others^31,32^ have established several frameworks for investigating the mating duration using sophisticated genetic techniques that can analyze and uncover the neural circuits’ principles governing interval timing. In particular, males exhibit LMD behavior when they are exposed to an environment with rivals, which means they prolong their mating duration. Conversely, they display SMD behavior when they are in a sexually saturated condition, meaning they reduce their mating duration^33,34^.

Multiple systems provide evidence that neuromodulators, such as biogenic amines or neuropeptides, are crucial internal state and behavior regulators^35–38^. Internal states prolong external influences on behavior^39^ and neuropeptides may facilitate internal state perpetuation^3^. SIFa is a conserved neuropeptide in insects, crustaceans, and arachnids. This neuropeptide is expressed in four cells near pars intercerebralis (PI) and the neuronal process of these neurons extend throughout the neural system^30,40–43^. Studies have linked SIFa to hunger and feeding^44,45^, courtship^41,42^, sleep^40,46,47^, and memory^48^. Here, we provide clear evidence that broadly arborized SIFa neurons record the internal states for two distinct interval timing behaviors.

## RESULTS

### The expression of SIFa in neurons controls two unique interval timing behaviors in male *Drosophila melanogaster*, regardless of sexual dimorphism

To determine which neuropeptide is neuronally connected to both LMD and SMD (Fig. 1A and 1B), we performed screening with pan-neuronal *elav^c^*^155^ driver with neuropeptide and neuropeptide receptor-RNAi lines (Table S1 and Document S1. Fig. S1–S5). We discovered that knocking down the neuronal expression of CCHa1, CCHa2, ETH, ilp4, and SIFa eliminates both LMD and SMD behaviors (Table S1 and Fig. 1C and 1D). SIFa is selected from the neuropeptides that affect both LMD and SMD because it is known to govern internal state-related activities such as sleep, hunger, and mating^40,42,44–46^. Glial knockdown of SIFa didn’t affect LMD and SMD (Fig. 1E-F) indicating that SIFa expression in glia is not associated with interval timing behaviors. The expression pattern of SIFa in four neurons located in PI is validated by utilizing glial-, neuronal-, or VNC-expressing *GAL80* lines (Fig. 1G; Fig. S1A, S1G and S1H), as well as anti-repo or anti-elav antibodies (Fig. S1B). Localized amplification of SIFa protein level through the utilization of *UAS-SIFa* with *GAL4^SIFa.PT^* influences both LMD and SMD behavior through either secretion (Fig. 1H and 1I) or membrane-tethered formats (Fig. 1J and 1K), suggesting that the level of SIFa protein in SIFa neurons is important for controlling interval timing behaviors. However, we were unable to identify any variations in the degree of *SIFa* expression across distinct internal states (Fig. 1L; Fig. S1O and S1Q). These findings indicate that both the neuronal activity and plasticity of SIFa neurons, as well as the SIFa neuropeptide itself, play a critical role in regulating interval timing behaviors.

**Fig. 1.**
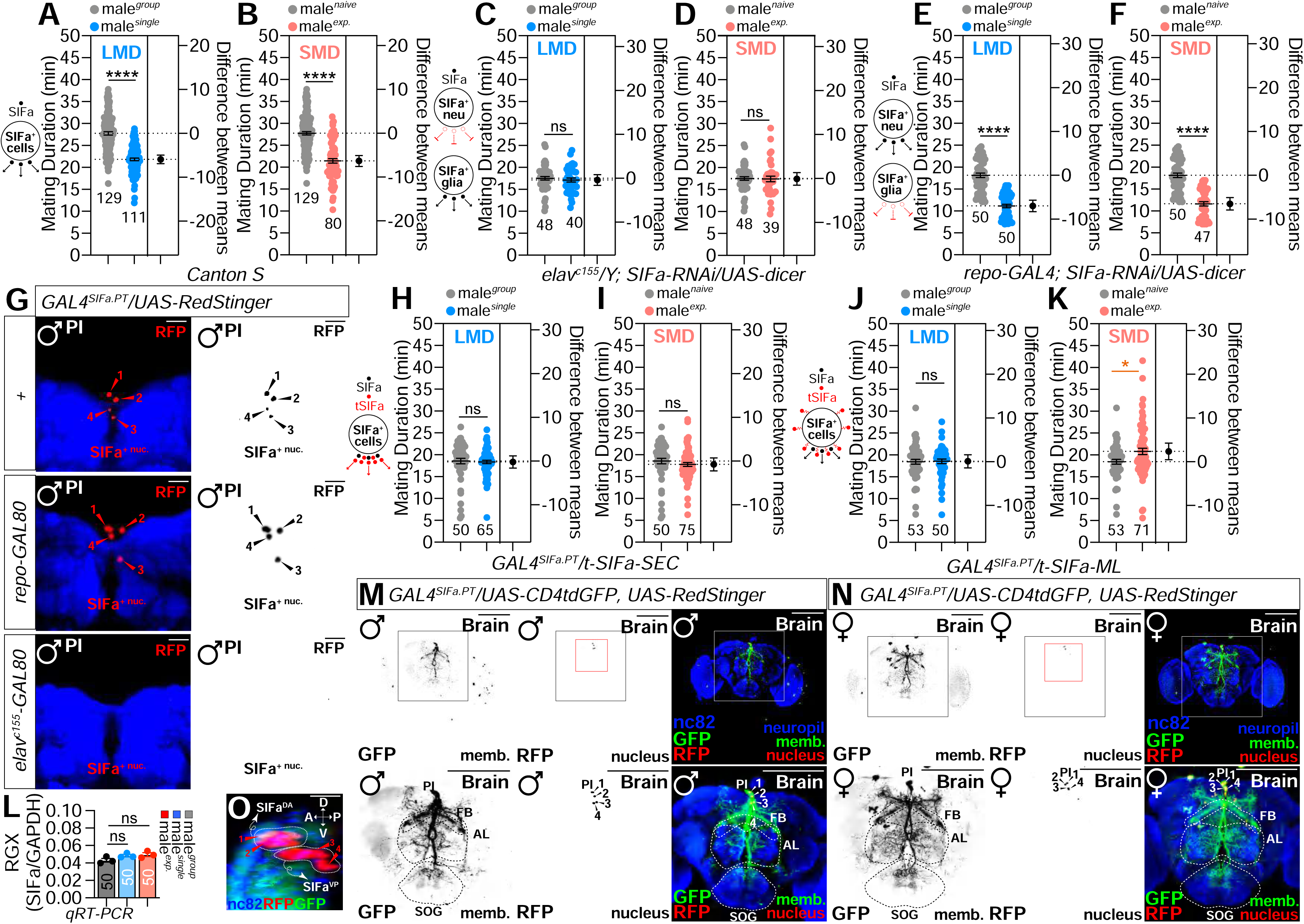
Neuronal expression of *SIFa* controls interval timing behaviors.(A) LMD assays of CS males. Light grey dots represent group males and blue dots represent single reared ones. Dot plots represent the MD of each male fly. The mean value and standard error are labeled within the dot plot (black lines). Asterisks represent significantdifferences, as revealed by the Student’s t test (* p<0.05, ** p<0.01, *** p<0.001). The same notations for statistical significance are used in other figures. (B) SMD assays of CS males. Light grey dots represent naïve males and pink dots represent experienced ones. (C-F) Mating duration (MD) assays for *GAL4* mediated knockdown of SIFa via *SIFa-RNAi* using the *elav^c^*^155^ driver (C-D), and *repo-GAL4* driver (E-F). (G) Flies expressing *GAL4^SIFa.PT^/+*(top) or *repo-GAL80; GAL4^SIFa.PT^* (middle) or *elav-GAL80; GAL4^SIFa.PT^* drivers(bottom) together with *UAS-RedStinger* were immunostained with anti-SIFa (red) antibodies. Red arrowheads indicate SIFa region labeled by anti-SIFa antibodies. The right panels are presented as a grey scale to clearly show the axon expression patterns of *SIFa* neurons in the adult brain labeled by *GAL4* drivers. Scale bars represent 10 µm. (H-K) MD assays for GAL4-driven over expression of *SIFa* via *UAS-t-SIFa-SEC* and *UAS-t-SIFa-ML*. (L) qRT-PCR results show that SIFa expression level is not affected by group, single and exp conditions. See the MATERIALS AND METHODS for a detailed description of the Quantitative RT-PCR used in this study. (M-N) Male (left) and female (right) flies expressing *GAL4^SIFa.PT^* together with *UAS-CD4tdGFP* and *UAS-RedStinger* were immunostained with anti-GFP (green), anti-RFP (red), nc82 (blue) antibodies. areas outlined by white boxes are enlarged in the bottom panel. Scale bars represent 100 µm. (O) *SIFa* neurons are categorized into two subpopulations, the anterior-dorsal *SIFa* neurons (SIFa^DA^) and the posterior-ventral *SIFa* neurons (SIFa^VP^), based on their anatomical positioning and putative functional roles. D: dorsal; V: ventral; A: anterior; P: posterior. Scale bars represent 10 µm. See also Fig. S1. and movie 1.

In order to compare the distinctions between male and female SIFa-positive neurons in the central nervous system, we examined the distribution of SIFa-expressing cells in terms of their membrane and nuclear expression patterns, utilizing *UAS-CD4-tdGFP* and *UAS-RedStinger*. As previously described^43,49^, a membrane marker expressing *GAL4^SIFa.PT^* exhibits a broad arborization pattern in both the male and female central brain (Fig. 1M and 1N). We found no significant differences between male and female brain regions covered by the SIFa cell membrane (Fig. S1C to S1E). SIFa acts on *fruitless*-positive neurons via the SIFa receptor (SIFaR) and it has been reported that SIFa neurons do not express fruitless, which are responsible for the generation of male-specific neuronal circuitry^41^. We have validated that *GAL4^SIFa.PT^*neurons do not label *fru^FLP^*-positive neurons in the male brain (Fig. S1F). Interval timing behaviors are unaffected by the expression of female form of doublesex (*UAS-dsx^F^*) (Fig. S1I and S1J), transformer (*UAS-tra^F^*) (Fig. S1K and S1L), or sex lethal (*UAS-sxl*) (Fig. S1M and S1N) all of which have strong feminizing activity. According to these results, neurons that express SIFa are not implicated in sexual dimorphism by itself.

### Architecture of SIFa-positive neurons in the central nervous system

Four SIFa neurons located in the PI are likely the most extensively branching peptidergic neurons in *Drosophila*^50^. We used an interactive tool for neuron connectivity (Virtual Fly Brain)^51^ to precisely determine the location of SIFa-positive neurons. We discovered that the cell bodies of SIFa neurons arborize from the posterior to the anterior region (Fig. S1E). It was verified through 3D reconstruction that two dorsal cell bodies are situated further anteriorly than two ventral cell bodies (Mov. 1). Anterior-dorsal SIFa neurons (SIFa^DA^)and posterior-ventral SIFa neurons (SIFa^VP^) could potentially have distinct functions (Fig. 1O).

SIFa neurons exhibit extensive neuronal arborization and are recognized for their role in global control through a gradual and sustained mechanism^50^. Our analysis revealed that SIFa neurons display neuronal processes which cover the entire brain, regardless of sexes (Fig. 2A to 2C). The neuronal processes extend throughout approximately 40% of the brain, including the VNC, central brain (CB), and optic lobe (OL) in both males and females (Fig. 2D to 2I). By comparison, SIFa neuronal processes cover only 10% of the larval CNS (Fig. 2J to 2L).

**Fig. 2.**
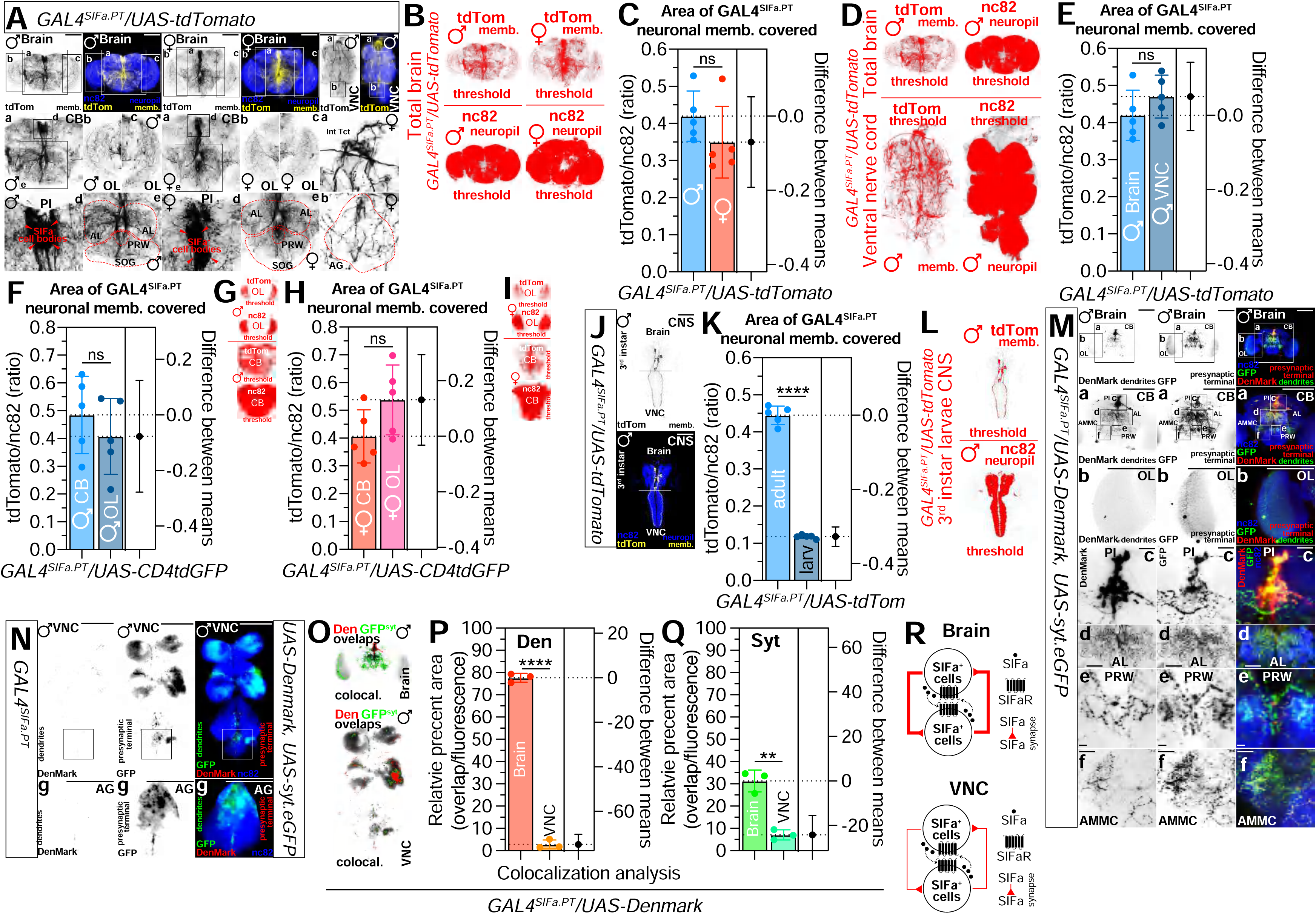
*SIFa* neurons display neuronal processes which cover the entire brain, regardless of sexes. (A) Flies expressing *GAL4^SIFa.PT^* drivers together with *UAS-tdTomato* were immunostained with anti-RFP (yellow), and nc82 (blue) antibodies. areas outlined by boxes are enlarged in the bottom panel, respectively. Arrowheads: cell body locations. Scale bars represent 100 µm. (B-C) Quantification of RFP fluorescence in the male (left) and female (right) fly brain expressing *GAL4^SIFa.PT^* driver together with *UAS-tdTomato*. (B) The threshold of RFP fluorescence (upper panel), nc82 (bottom panel) in male and female fly brain was marked by threshold function of ImageJ. (C) Quantification of RFP fluorescence in male and female brain. Bars represent the mean RFP fluorescence level with error bars representing SEM. Asterisks represent significant differences, as revealed by the Student’s t test and ns represents non-significant difference (*p<0.05, **p<0.01, ***p< 0.001). The same symbols for statistical significance are used in all other Figures. See the MATERIALS AND METHODS for a detailed description of the colocalization analysis used in this study. (D-E) Quantification of RFP fluorescence in brain and VNC of male fly. (F-I) Quantification of RFP fluorescence of CB and OL area in male and female. (J) Larva expressing *GAL4^SIFa.PT^*drivers together with *UAS-tdTomato* were immunostained with anti-RFP (yellow), and nc82 (blue) antibodies. Scale bars represent 100 µm. (K) Quantification of the RFP fluorescence in third instar larvae and adult. (L) The threshold of RFP fluorescence (top panel), nc82 (bottom panel) in male larva was marked by threshold function of ImageJ. (M-N) Distribution of dendrites and pre-synaptic terminals of neurons labeled via *GAL4^SIFa.PT^* in the brain and VNC. Flies expressing *GAL4^SIFa.PT^* together with *UAS-Denmark, UAS-syt.eGFP* were immunostained with anti-GFP (green), anti-DsRed (red) and nc82 (blue) antibodies. Areas outlined by boxes are enlarged in bottom panel. Scale bars represent 100 μm in M and 50 μm of N. (O-Q) Quantification of dendritic and presynaptic terminals of neurons labeled via *GAL4^SIFa.PT^* in the brain and VNC. (R) Schematic shows that SIFa+ cells form extensive synapses each other within the brain but not in the VNC. See also Fig. S2.

The connection between the dendrites and axons of the SIFa neuronal processes is unknown. To identify the dendritic and axonal components of SIFa-neuronal processes, we co-expressed *UAS-Denmark* (a dendritic marker) with *UAS-syt.eGFP* (a presynaptic marker) to label postsynapses and presynapses of neurons labeled by *GAL4*^52^. We discovered that dendrites of SIFa neurons span just the central brain area, including the PI, antennal lobe (AL), and posterior-ventral region of the protocerebrum (PRW), whereas presynaptic terminals cover nearly the whole brain region (magnified regions in Fig. 2M). Colocalization analysis in the total brain (Fig. S2A to S2C), the CB (Fig. S2D to S2F), and the OL (Fig. S2G to S2I) confirms the broader arborization of presynaptic terminals than dendrites.

Within the VNC, the majority of branching structures are found in the abdominal ganglion (AG) (Fig. 2N). Additionally, most of these branching structures are presynaptic terminals, which are different from those found in the brain (Fig. 2O to 2Q). This suggests that the majority of SIFa^+^ neuronal structures in VNC are axons rather than dendrites. Compared to brain SIFa^+^ dendrites, which overlap presynaptic terminals by almost 80%, VNC SIFa^+^ dendrites only overlap presynaptic terminals by around 2%. (Fig. 2P), suggesting that SIFa^+^ cells form extensive synapses each other within the brain but not in the VNC (Fig. 2R). We noticed that the *SIFa^2A-GAL^*^4^ strains, which were created through the knock-in (KI) lines using the site-specific integration system, exhibit robust expression exclusively in the brain region (Fig. S2J). This indicates the potential existence of distinct arborization and functional differences between the dorsal and ventral SIFa neurons (Fig. 1O), which may play a role in modulating physiology and behaviors.

### Glutaminergic and dopaminergic presynaptic inputs from SIFa-positive neurons influence interval timing behavior in a distinct manner from sleep behavior

It has been proposed that SIFa neurons might be able to utilize glutamate, a neurotransmitter, for controlling sleep^46^; however, no comprehensive analysis of neurotransmitter screening has been conducted yet. To determine which neurotransmitter (NT) system was responsible for the LMD/SMD, we utilized RNAi-mediated knockdown of key genes which are necessary to produce each NT. We chose *VGlut-RNAi* (*Vesicular glutamate transporter*) to disrupt the glutaminergic system (Fig. 3A and 3B), *ple-RNAi* (*pale*) for the dopaminergic system (Fig. 3C and 3D), *Gad1-RNAi* (*Glutamic acid decarboxylase 1*) for the GABAergic system (Fig. 3E and 3F), *Trhn-RNAi* (*Tryptophan hydroxylase neuronal*) for the serotonergic system (Fig. 3G and 3H), *Tdc2-RNAi* (*Tyrosine decarboxylase 2*) for the octopaminergic system (Fig. 3I and 3J), *ChAT-RNAi* (*Choline acetyltransferase*) for the cholinergic system (Fig. 3K and 3L), and *Hdc-RNAi* (*Histidine decarboxylase*) for the histaminergic system (Fig. 3M and 3N). We also conducted experiments on *dimm-RNAi*, a well-known transcription factor that plays a crucial role in the activity of neuroendocrine cells (Fig. S3A and S3B)^53^.

**Fig. 3.**
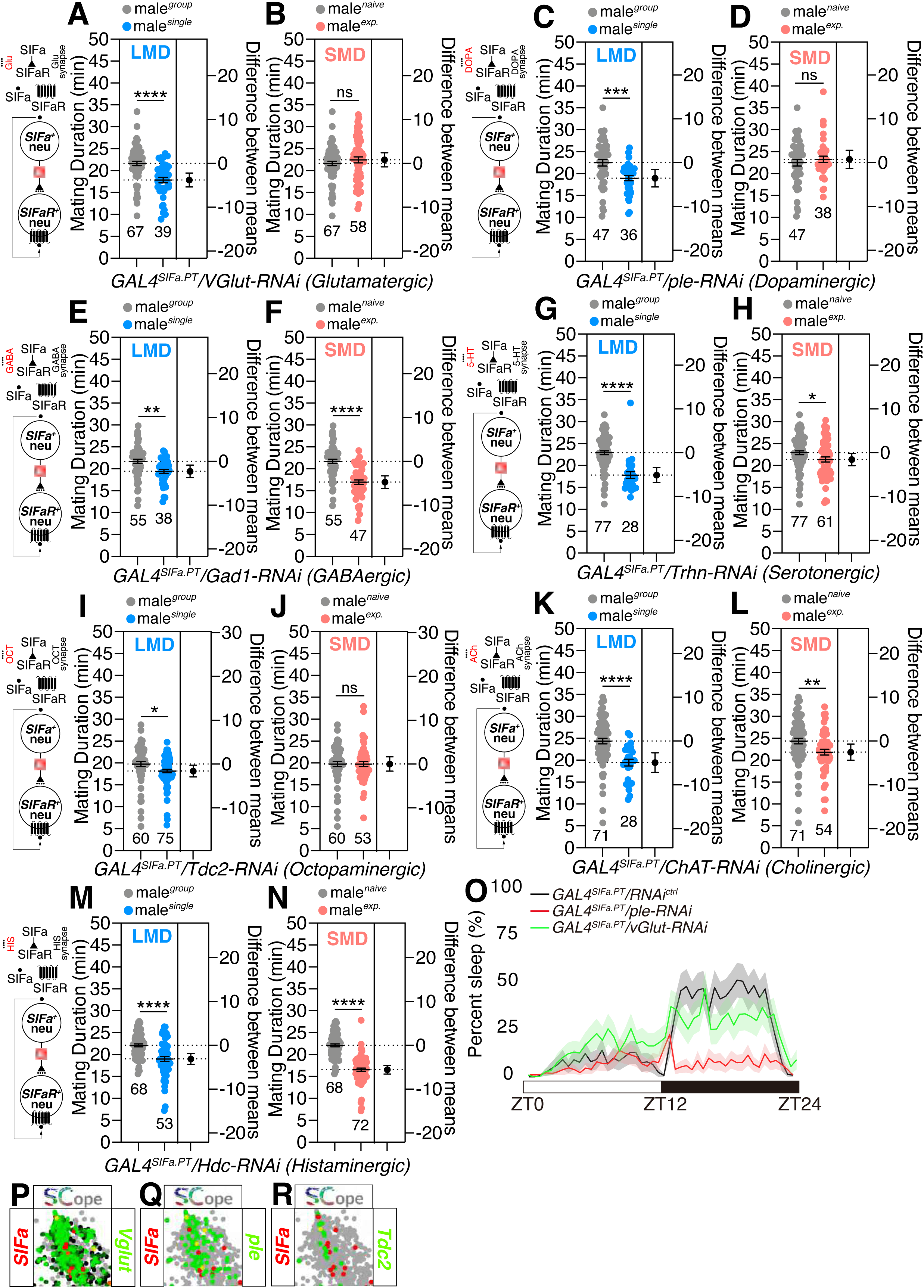
Interval time is affected by gluttaminergic and dopaminergic presynaptic inputs from SIFa-positive neurons. (A-N) MD assays of flies expressing the *GAL4^SIFa.PT^* driver together with (A-B) *VGlut-RNAi* (C-D) *ple-RNAi* (E-F) *Gad1-RNAi* (G-H) *Trhn-RNAi* (I-J) *Tdc2-RNAi* (K-L) *ChAT-RNAi* (M-N) *Hdc-RNAi*. The diagram illustrating the correlation between SIFa neurons, SIFaR, and the neurotransmitter is located on the left side. (O) Group sleep of flies expressing *GAL4^SIFa.PT^*driver together with *ple-RNAi* (red) less than heterozygous controls (grey) during the course of a 24-h day, but flies together with *VGlut-RNAi* (green) are not. (n = 100 flies per group). White and black bars denote periods of light and darkness. (P-R) Fly SCope single-cell RNA sequencing data of cells co-expressing *SIFa* together with *VGlut*, *ple* and *Tdc2*. See the MATERIALS AND METHODS for a detailed description of the Single-nucleus RNA-sequencing analyses—data and code availability used in this study. See also Fig. S3.

By using RNAi-mediated knockdown experiments, we found that glutaminergic, dopaminergic, and octopaminergic system are crucial for generating SMD but not LMD behavior in SIFa-positive neurons (Fig. 3A to 3N and Fig. S3A and S3B). SIFa also modulates the courtship behavior of male flies^49^. Studies have shown that social isolation can reduce male courtship behavior in *Drosophila melanogaster*^54–56^, however, courtship behavior does not change in response to sexual experience (Fig. S3C and S3D). The knockdown of VGlut or Tdc2 augmented the courtship activity of males reared in isolation, while diminishing it in males with prior sexual experience (Fig. S3E to S3H). The removal of ple, on the other hand, enhances courtship behavior in both socially isolated and sexually experienced males (Fig. S3G and S3H).

Notably, the sleep patterns and overall sleep duration of male flies were not influenced by the interruption of glutaminergic synaptic transmission from SIFa neurons (Fig. 3O). The findings indicates that glutaminergic inputs specifically influence SMD behavior, but do not have an effect on sleep. On the other hand, dopaminergic inputs from SIFa neurons have an impact on both sleep and SMD behavior. The Fly SCope single-cell RNA sequencing data indicates that SIFa neurons exhibit the highest level of co-expression for most neurotransmitters, with the exception of Trhn and Hdc (as shown in Fig. 3P to 3R; Fig. S3K to S3P). All these data indicate that the modulation of diverse behavioral repertoires can be achieved by the use of various combinations of neurotransmitter co-transmission in SIFa neurons.

### sNPF inputs are critical to generate SIFa-mediated internal states for interval timing behavior

The short neuropeptide F (sNPF) is involved in executive signaling inside local neuroendocrine cells. It functions as a co-transmitter through its receptor, sNPF-R, in specific circuits. The sNPF governs the regulation of food intake and body size^57,58^, is sleep-promoting inhibitory modulator^59–61^, affect olfactory memory formation^62^, and control interval timing behavior^33^. The idea proposes that sNPF signaling transmits hunger and satiety information to SIFa neurons in order to control food search and feeding^44^. However, there is currently no reported direct evidence to support this hypothesis.

Knockdown of sNPF in SIFa-expressing neurons (Fig. 4A and 4B) or SIFa in sNPF-expressing cells (Fig. 4C and 4D) has no impact on both LMD and SMD behavior, implying that both neuropeptide-expressing cells are mutually exclusive. Knockdown of sNPF-R in SIFa-expressing cells (Fig. 4E and 4F) or SIFa in sNPF-R-expressing cells (Fig. 4G and 4H) disrupts only SMD behavior, implying that sNPF-R function in SIFa-positive cells and SIFa function in sNPF-R-positive cells are required to generate SMD. SIFa transmits information to adjacent cells through the SIFa Receptor (SIFaR). Inhibition of SIFaR in cells expressing sNPF leads to the disruption of both LMD and SMD behaviors (Fig. 4I and 4J). Similarly, the knockdown of sNPF in cells expressing SIFaR also disrupts both behaviors (Fig. 4K and 4L). When the expression of SIFaR was suppressed in cells expressing sNPF-R using *sNPF-R^64H^*^09^*-GAL4* drivers, only the SMD behavior was affected (Fig. S4A and S4B). However, when the expression of sNPF-R was suppressed in SIFaR-expressing cells, both LMD and SMD behaviors were disrupted (Fig. S4C and S4D). These data indicate that the signaling of SIFa-SIFaR is regulated by sNPF-sNPF-R signaling, and that SIFa and sNPF collaborate to regulate interval timing behaviors in a specific manner.

**Fig. 4.**
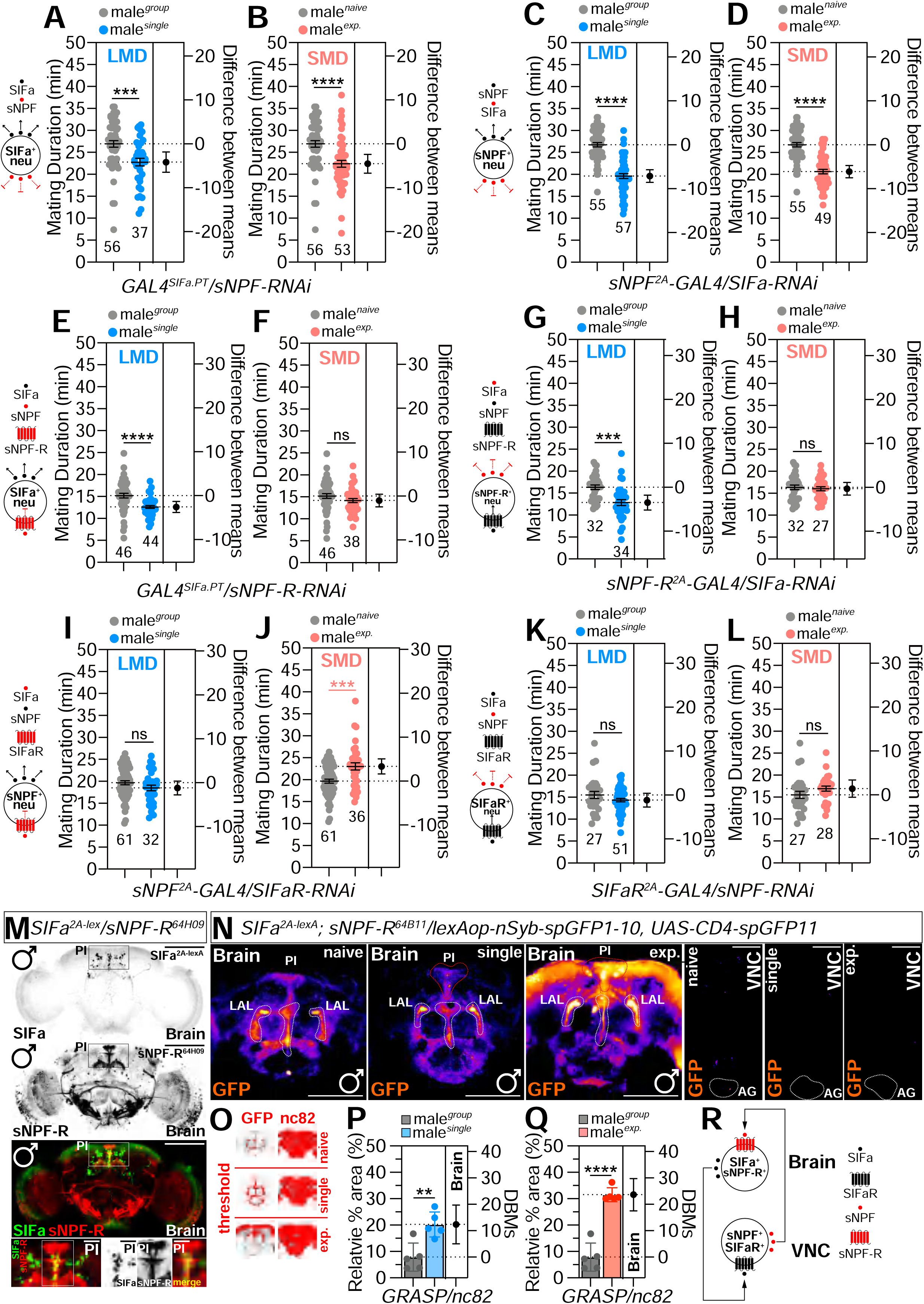
*sNPF* signals are necessary to create internal states mediated by *SIFa*. (A-B) MD assays of flies expressing the *GAL4^SIFa.PT^* driver together with *sNPF-RNAi*. (C-D) MD assays of flies expressing the *sNPF-GAL4* driver together with *SIFa-RNAi*. (E-F) MD assays of flies expressing the *GAL4^SIFa.PT^* driver together with *sNPF-R-RNAi*. (G-H) MD assays of flies expressing the *sNPF-R^2A^-GAL4* driver together with *SIFa-RNAi*. (I-J) MD assays of flies expressing the *sNPF^2A^-GAL4* driver together with *SIFaR-RNAi*. (K-L) MD assays of flies expressing the *SIFaR^2A^-GAL4* driver together with *sNPF-RNAi*. (M) male flies expressing *SIFa^2A-lexA^* and *sNPF-R^64H^*^09^*-GAL4* drivers together with *lexAop-mCD8RFP* and *UAS-mCD8RFP* were imaged live under a fluorescent microscope with anti-GFP (green), anti-DsRed (red), and nc82 (blue) antibodies. yellow arrows indicate *SIFa*-positive neurons and *sNPF-R*-positive neurons. Scale bars represent 100 μm. (N) GRASP assay for *SIFa^2A-lexA^* and *sNPF-R^64B^*^11^*-GAL4* in male brain (left three columns) and VNC (right three columns). Male flies expressing *SIFa^2A-lexA^.* , *sNPF-R^64B^*^11^*-GAL4* and *lexAop-nsyb-spGFP1-10, UAS-CD4-spGFP11* were dissected after 5 days of growth (mated male flies had 1-day of sexual experience with virgin females). The white dashed line highlights the GRASP signal. Scale bars represent 100 μm. (O-Q) Quantification of synapses formed between *SIFa^2A-lexA^* and *sNPF-R^64H^*^09^*-GAL4* in the brain. (O) The threshold of GFP fluorescence (left panel), nc82 (right panel) of naïve, single and exp in male fly brain was marked by threshold function of ImageJ. (P-Q) Quantification of synapses formed between *SIFa^2A-lexA^* and *sNPF-R^64H^*^09^*-GAL4* in the brain. Revealed by the GRASP system in naïve, single and mated male flies. GFP fluorescence was normalized to that in nc82. See the MATERIALS AND METHODS for a detailed description of the Quantitative analysis of fluorescence intensity used in this study. (R) Schematic depicting the interaction between *SIFa* and *sNPF* neurons. See also Fig. S4.

The coexpression of SIFa with sNPF-R was verified by genetic intersection approaches (Fig. 4M; Fig. S4E) and fly SCope data (Fig. S4H and S4I). The extensive coexpression patterns of sNPF with SIFaR were also validated using the same methodology (Fig. S4F and S4G; Fig. S4J to S4L). The expression of SIFaR and sNPF-R is highly prevalent in neuronal populations together with sNPF (Fig. S4M and S4N). Upon performing GRASP using *SIFa^2A-lexA^*in conjunction with *sNPF-R^64B^*^11^*-GAL4*, we observed the formation of robust synapses in the PI tract and the lateral accessory lobe (LAL) area connecting them. Interestingly, when males are exposed to social isolation or sexual experience, there is an increase in synaptic signals around the PI region where SIFa cells are located (Fig. 4N). This indicates that the synapses between SIFa cells expressing sNPF-R become stronger (Fig. 4O to 4Q). In contrast, we found no synapses between *SIFa^2A^* and *sNPF-R^64H^*^09^ in VNC (Fig. 4N). These findings demonstrate that the synaptic plasticity among SIFa cells in the brain plays a crucial role in regulating interval timing behaviors.

The cells that express both sNPF and SIFaR are predominantly found in the AG (shown by yellow dotted lines labeled as "cell" in Fig. S4G). This indicates that the sNPF-positive cells responsible for SIFa-SIFaR signaling to induce SMD are located in the VNC rather than the brain. The results suggests that SIFa signals originating from the brain and sNPF signals originating from the VNC create a feedback or feedforward loop to modulate the internal states of the central nervous system (CNS) in male flies when they are exposed to varying social environments. The internal states created by the interaction of these two neuropeptides and their receptors, which signal through the brain to VNC, are crucial components of the circuit responsible for interval timing behaviors (Fig. 4R).

### Neuropeptides that communicate to SIFa neurons exert a distinct influence on the interval timing behaviors

The SIFa neurons receive inputs from many peptidergic pathways including Crz, dilp2, Dsk, sNPF, MIP, and hugin, however this speculation is based on morphological observations^44,50,63^. In order to determine which inputs are essential for interval timing behavior, we systematically examined the role of each neuropeptide receptor in SIFa-positive cells. When we reduced the CrzR in SIFa cells, both LMD and SMD remained unaffected, indicating that Crz signaling is not necessary for SIFa-mediated interval timing behavior (Fig. 5A and 5B). Hugin and MIP modulates satiety and hunger via SIFa neurons^44^. The inhibition of hugin receptor, PK2-R2, or MIP receptor, CCHa1R, did not have an impact on LMD or SMD behavior. This indicates that the transmission of signals through MIP and hugin peptides to SIFa neurons does not influence interval timing behavior (Fig. 5C to 5F).

**Fig. 5.**
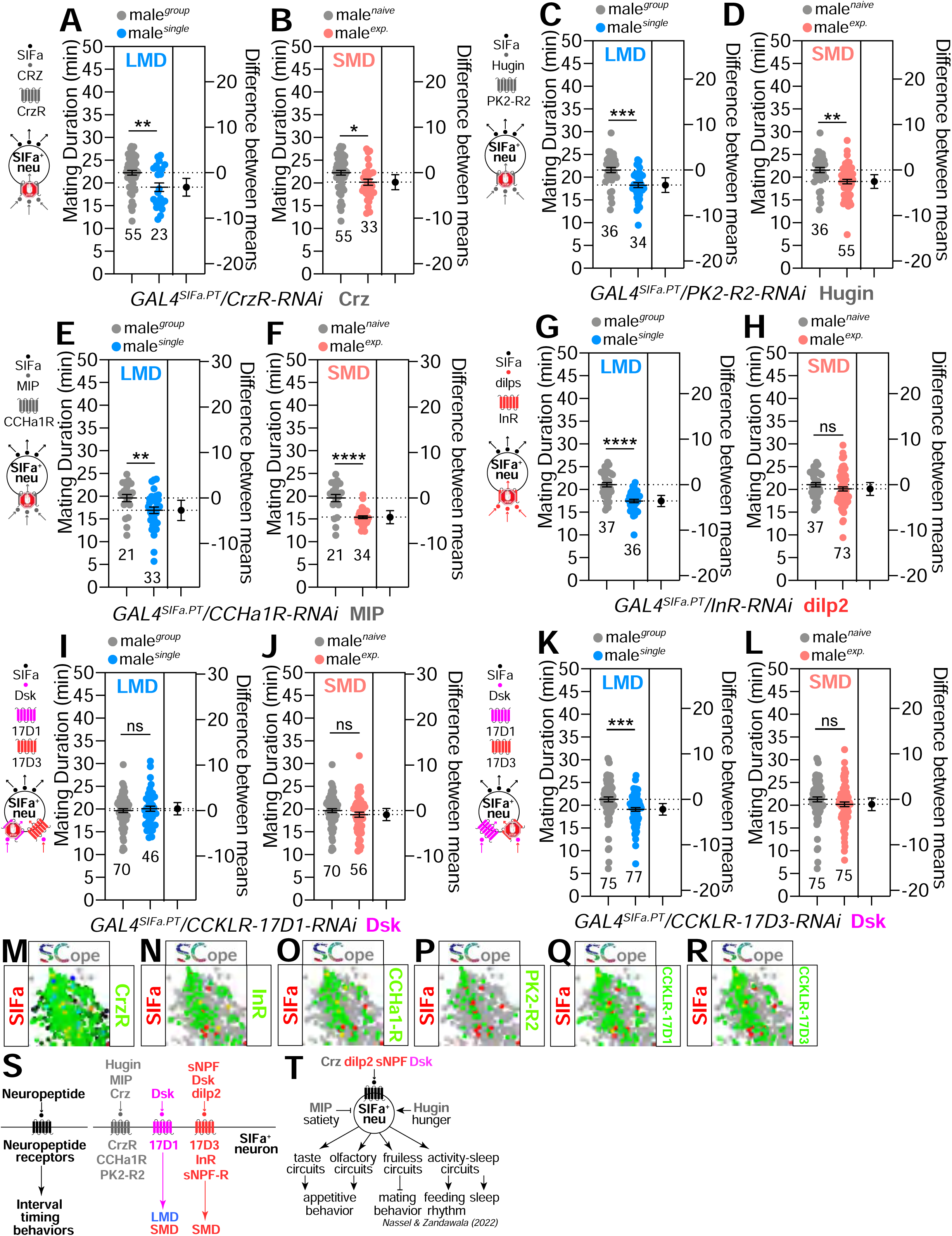
Neuropeptides interacting with *SIFa* neurons are implicated in the modulation of interval timing. (A-L) MD assays of flies expressing the *GAL4^SIFa.PT^* driver together with (A-B) *CrzR-RNAi* (C-D) *PK2-R2-RNAi* (E-F) *CCHa1R-RNAi* (G-H) *InR-RNAi* (I-J) *CCKLR-17D1-RNAi* (K-L) *CCKLR-17D3-RNAi*. The diagram of the relationship between *SIFa* neurons and neuropeptide receptors is on the left. (M-R) Fly SCope single-cell RNA sequencing data of cells co-expressing *SIFa* together with *CrzR*(M), *InR*(N), *CCHa1-R*(O), *PK2-R2*(P), *CCKLR-17D1*(Q), and *CCKLR17D3*(R). (S) Illustration depicting the influence of neuropeptide receptors expressed on *SIFa* cells. (T) Schematic representation of the modulatory effects of *SIFa* neuropeptides on diverse behaviors. See also Fig. S5.

The inhibition of InR in SIFa cells exclusively impacts SMD behavior, indicating that dilp2-InR signaling in SIFa neurons is solely necessary for SMD behavior (Fig. 5G and 5H). Dsk-expressing neurons are a type of interneuron that play a crucial role in orchestrating various signals related to satiety^64^, aggression^65^, courtship, daytime locomotor activity, gustatory reception, and feeding^66^. DSK employs two receptors that are functionally different, namely CCKLR-17D1 and CCKLR-17D3^65,67–69^. The inhibition of CCKLR-17D1 in SIFa neurons disrupted both LMD and SMD behaviors (Fig. 5I and 5J), whereas CCKLR-17D3 only affected SMD behavior (Fig. 5K and 5L). This indicates that Dsk can selectively regulate LMD and SMD behaviors through its functionally distinctive receptors. Except for PK2-R2, all other genes coexpress with SIFa in SCope data, indicating that hugin inputs to SIFa may not be transmitted through peptidergic signaling. Therefore, out of the many peptidergic inputs to SIFa, only Dsk, sNPF, and dilp2 have a particular regulatory effect on either LMD or SMD behavior (Fig. 5S). This is particularly intriguing given that courtship behavior, which is known to be regulated by SIFa neurons^41,49^, is influenced by all of the peptidergic inputs described above (Fig. S5A to S5N). However, it appears that various peptidergic inputs to SIFa neurons influence courtship behavior that is under context-dependent modulation (Fig. S5O). The data indicate that various peptidergic inputs to SIFa influence interval timing and courtship behavior in an entirely distinct manner (Fig. 5T).

### The neuronal activity and synaptic plasticity of SIFa neurons determine internal states for interval timing behavior

The activity of SIFa neurons can be changed by a number of different stimuli. However, the precise mechanisms by which they regulate SIFa neuronal activity remain unknown. To determine whether neuronal activities undergo alterations in SIFa neurons associated with interval timing, we utilized the TRIC (transcriptional reporters for sensing Ca^2+^) system^70^. TRIC serves as a valuable complement to functional Ca^2+^ imaging by integrating long-term changes in neuronal activity and providing genetic access to neurons based on their activity. Given that both LMD and SMD require at least 6-12 hours of social isolation or sexual interaction^71,72^, repeated sensory inputs could potentially lead to the accumulation of the modified transcription factor within the nucleus of activated neurons *in vivo*. Indeed, social isolation or sexual experiences affected the neural activity of *GAL4^SIFa-PT^*-labeled neurons (Fig. 6A). Male flies with social isolation or sexual experience exhibited robust TRIC fluorescence in the PI region (Fig. 6B and 6C). In contrast, the TRIC fluorescence in the prow (PRW) region becomes weakened in males raised in isolation and stronger in males with sexual experience (Fig. 6D and 6E). Considering the similarity of the control RFP signals across conditions, we conclude that social isolation and sexual experience alter Ca^2+^ levels in SIFa neurons in a comparable yet distinct manner (Fig. S6-1A).

**Fig. 6.**
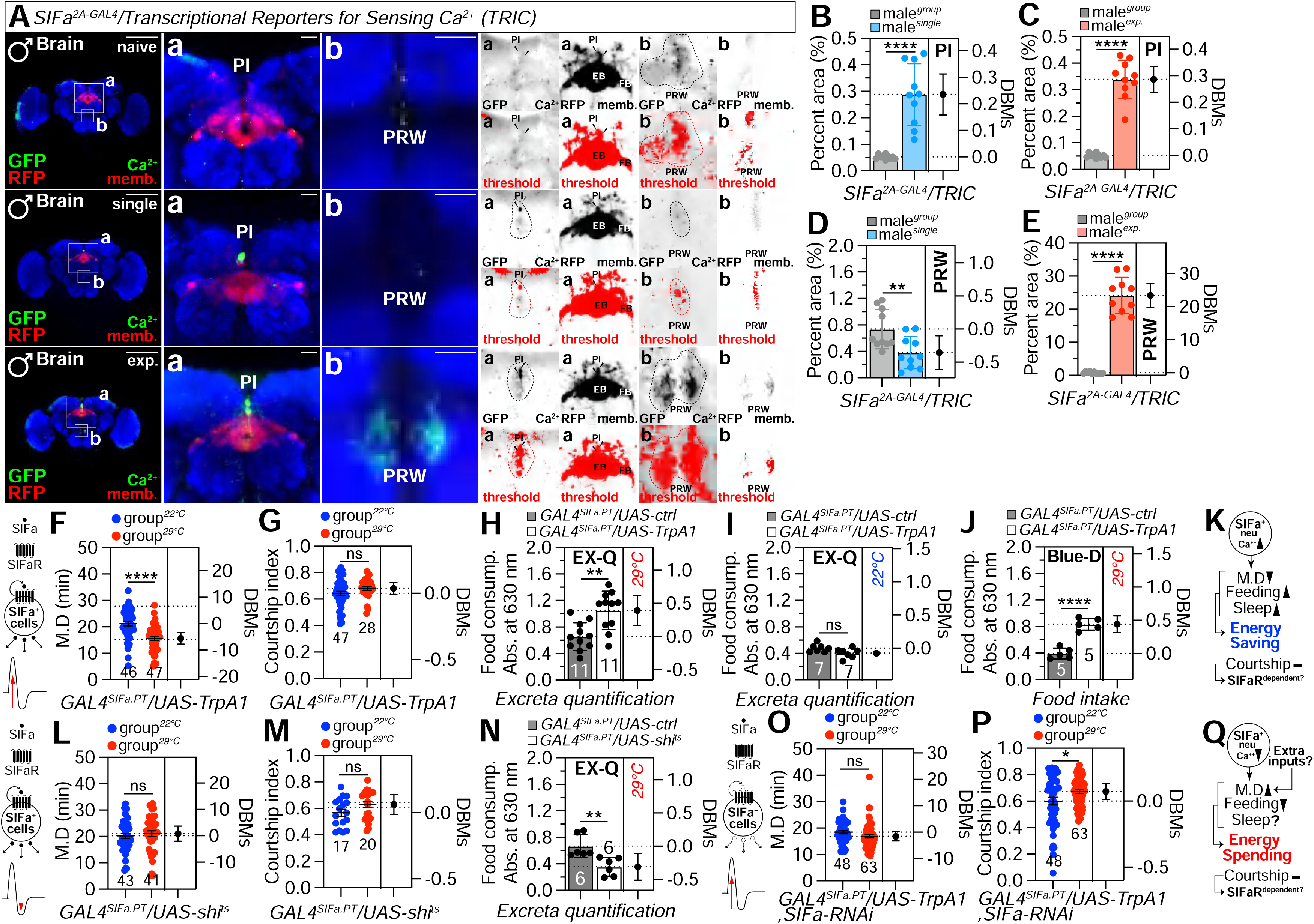
Internal interval timing behavior is determined by *SIFa* neuronal activity and synaptic plasticity. (A) CaM/MKII-mediated TRIC signal in the brain of transgenic flies (*nsyb-MKII::GAL4DBDo, UAS-p65AD::CaM, 10XUAS-IVS-mCD8::RFP, 13XLexAop2-mCD8::GFP*). The right four panels are presented as a gray scale to clearly show the TRIC signal of *SIFa* neurons in PI and PRW labeled by *SIFa^2A-GAL4^* driver and TRIC. Scale bars represent 100 μm in left panels and 10 μm in a and b. (B-E) Quantification of GFP signal of male fly expressing *SIFa^2A-GAL4^* drivers together with Transcriptional Reporters for Sensing Ga^2+^ of PI and PRW region. (F-G, L-M) Temperature-shift mating duration and courtship index of male fly expressing *GAL4^SIFa.PT^* drivers together with heat-sensitive *Drosophila* cation channel *TrpA1* and *shi^ts^*. (H-J) 24-h food intake of males measured by EX-Q and Blue-Dye assays of male flies expressing *GAL4^SIFa.PT^* driver together with *TrpA1* on yeast-sugar medium of different temperature (H, 29℃; I, 22℃; J, 29℃). See the MATERIALS AND METHODS for a detailed description of the EX-Q and Blue-Dye assay used in this study. (N) quantification of 24-h food intake of males expressing *GAL4^SIFa.PT^* driver together with shi^ts^ measured by EX-Q on yeast-sugar medium on 29℃. (O-P) Analysis of mating duration and courtship index in male *Drosophila* expressing *GAL4^SIFa.PT^* drivers, in conjunction with the heat-sensitive *Drosophila* cation channel *TrpA1* and *SIFa-RNAi*, following temperature shifts. (K and Q) *SIFa* neurons modulate energy expenditure through the fine-tuning of their intracellular environments. See also Fig. S6.

Next, to determine if the temporal activation of SIFa neurons may generate distinct internal states for mating duration in the absence of social isolation of sexual experience, we expressed the heat-sensitive *Drosophila* cation channel TrpA1 in SIFa neurons and then transferred the experimental group to the activation temperature (29°C). Surprisingly, flies expressing TrpA1 in SIFa neurons at the activation temperature exhibited a shorter mating duration than flies maintained at 22°C (Fig. 6F) compared to genetic control (Fig. S6-1B and S6-1C). However, the temporal activation of SIFa neurons did not have an impact on courtship behavior (Fig. 6G). This indicates that the mating duration and the level of courtship activity are controlled separately by the activity of SIFa neurons. Stimulation of SIFa neurons resulted in an elevation in food consumption, as assessed using the EX-Q (excreta quantification) method^73^ (Fig. 6H and 6I) and the colorimetric food intake assay^74^ (Fig. 6J and Fig. S6-1E). Notably, the temporary deactivation of SIFa neurons through the expression of shi^ts^ did not have any impact on the mating duration or the level of courtship activity (Fig. 6L to 6M; Fig. S6-1D) within the temperature range of 22°C to 29°C. Nevertheless, the temporary deactivation of SIFa neurons leads to a decrease in food consumption in male flies (Fig. 6N; Fig. S6-1F to S6-1H). These findings indicate that activating SIFa-positive neurons is sufficient to generate specific internal states that lead to reduced mating durations and increased food intake. However, it is not sufficient to alter courtship activities (Fig. 6K). Moreover, the deactivation of SIFa neurons has the potential of generating internal states that decrease food consumption, but it does not have the ability to alter the mating duration or courtship behavior (Fig. 6Q).

We then tried to alter the neuronal activity of SIFa-positive neurons during social isolation and sexual experiences by expressing various modifiers of neuronal activity with *GAL4^SIFa.PT^*. By expressing TNT (tetanus toxin light chain) to block synaptic transmission in SIFa neurons, we observed that both LMD and SMD were impaired (Fig. S6-1I and S6-1J). However, food intake was shown to be elevated under these circumstances (Fig. S6-1K and S6-1L). In contrast to TNT expression, potassium channels KCNJ2/OrkΔC^75^ or bacterial sodium channel NachBac expression in SIFa neurons only impaired either LMD or SMD (Fig. S6-1M and S6-1N; Fig. S6-1U and S6-1V; Fig. S6-1Q and S6-1R). Notably, the food intake was partially increased by constitutive modulation of SIFa neuronal activities whether neuronal activity was activated or inhibited (Fig. S6-1O and S6-1P; Fig. S6-1S and S6-1T). TNT has been used effectively in several studies to inhibit neurotransmitter and neuropeptide release, KCNJ2/OrkΔC prevents membrane depolarization, and NachBac induces sodium conductance, which causes cell depolarization^76^. Thus, our research suggests that the release of neurotransmitter and neuropeptide vesicles is necessary for the transition of internal states, leading to the production of both LMD and SMD, as well as regulating food intake (Fig. S6-1W). When TrpA1 artificially stimulated SIFa neurons while simultaneously knocking down SIFa, the mating duration remained same (Fig. 6O). However, there was a modest increase in courtship activity (Fig. 6P). The data supports our hypothesis that altering internal states for interval timing activities requires augmentation between SIFa synaptic transmission and SIFa release (Fig. S6-1W). Furthermore, the knockdown of Tdc2, but not VGlut or ple, abolished the reduced mating duration phenotype induced by artificial activation of SIFa neurons (Fig. S6-2X-Z), indicating that octopamine, rather than dopamine or glutamate, is the critical neurotransmitter involved in the induction of energy balance behaviors through SIFa co-transmission. This suggests that octopamine signaling, specifically, is essential for mediating the behavioral response to SIFa inputs in the context of mating duration regulation.

### Internal states of a socially isolated or sexually experienced male’s nervous system represent neuronal plasticity of SIFa neurons

Even though calcium activity successfully represents the internal states that determine the mating duration of male flies, it is still unknown why similar neuronal alteration affect LMD and SMD in different ways (Fig. S6-1M and S6-1N; Fig. S6-1Q and S6-1R; Fig. S6-1U and S6-1V). Thus, we used *DenMark* and *syt.eGFP* to measure the synaptic plasticity of SIFa neurons in various conditions due to their capacity to detect broad changes in synaptic structure. DenMark signals in the brains of grouped (naïve), singly reared (single), and sexually experienced (exp.) males were comparable; however, syt.eGFP signals in the brains of group reared males were significantly stronger than those in the brains of both singly reared and sexually experienced males (Fig. 7A to 7E). Similar to the brain, the syt.eGFP signals in the VNC of singly reared and sexually experienced males were significantly weaker than in the VNC of grouped males (Fig. S7A to S7E). The findings indicate that significant alterations in the postsynaptic region, rather than the presynaptic region, occur in the central nervous system of males who are in groups, socially isolated, or undergo sexual experience (Fig. 7F). These variations in extensive structural synaptic changes may explain why various neuronal activity alterations influence LMD and SMD in different ways.

**Fig. 7.**
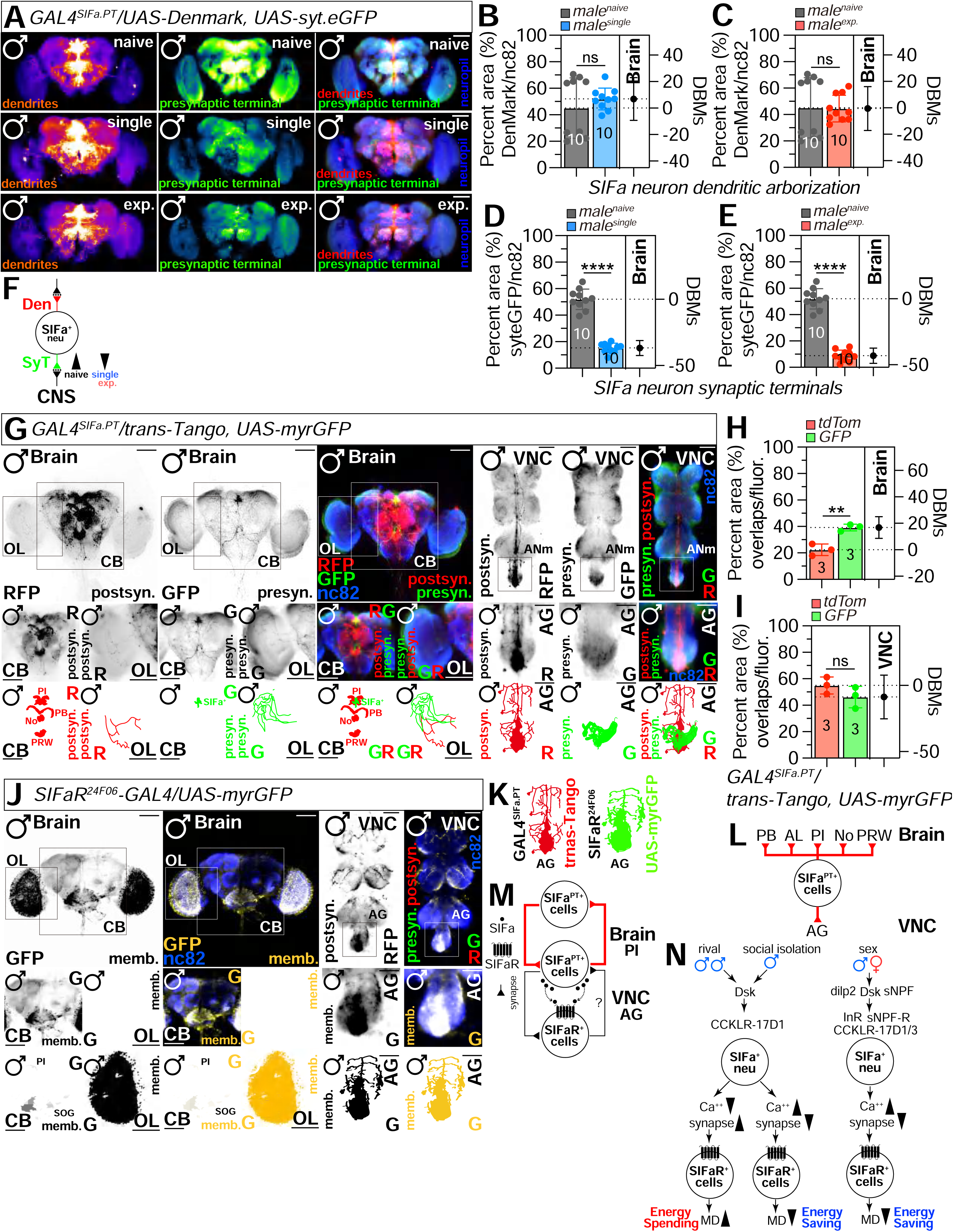
*SIFa* neuronal plasticity is seen in socially isolated or sexually experienced males’ neural systems. (A) image of a brain in which *GAL4^SIFa.PT^* was used to simultaneously drive expression of the dendritically-localized Denmark (red) and presynaptic terminal (green) of naïve (top), single (middle) and mated (bottom) male flies. Scale bars represent 100 μm. (B-E) Quantification of dendritic arborization and synaptic terminals of *SIFa* neurons in Fig. A. (F) Schematic representation of dendritic arborization and presynaptic terminal dynamics in *SIFa* neurons subjected to various experimental conditions. (G) Confocal image of male brain and VNC in which *GAL4^SIFa.PT^* was used to simultaneously drive expression of trans-Tango(magenta), visualized with RFP antibody staining, which identifies postsynaptic cells, and myrGFP (green), which labels the *GAL4^SIFa.PT^* cells. Areas outlined by black boxes in top panels are enlarged in middle. The bottom panels are presented as a red scale to show the threshold of RFP and GFP signals marked by threshold function of ImageJ. Scale bars represent 100 μm. (H-I) Quantification of GFP and RFP fluorescence in male fly brain (H) and VNC (I) expressing *GAL4^SIFa.PT^* together with *UAS-myrGFP* and *UAS-trans-Tango*. (J) male flies expressing SIFaR24F06-GAL4 drivers together with UAS-myrGFP were imaged live under a fluorescent microscope with anti-GFP (yellow), and nc82 (blue) antibodies. areas outlined by boxes are enlarged in the bottom panel. The bottom panels are presented as a yellow scale to show the threshold of GFP signals marked by threshold function of ImageJ. Scale bars represent 100 μm in brain and CB, 10 μm in OL. (K) Comparative examination of AG region SIFa trans-Tango fluorescence signals and SIFaR expression. (L) Schematic illustration of *SIFa* neurons projecting to PB, AL, AI, PRW, AG etc. to regulate different behaviors. (M) Schematic depiction of the anatomical arrangement and functional connectivity between *SIFa* neurons located in the PI of the brain and *SIFaR* neurons situated in the AG of the VNC, demonstrating their coordinated involvement in *SIFa-SIFaR* signaling. (N) Schematic showing how neuropeptide receptor activation regulates energy conservation and spending in Drosophila melanogaster. See also Fig. S7 and S8.

Finally, we sought to identify circuits downstream of *GAL4^SIFa.PT^* neurons. At first, we used a recently developed system for unbiased trans-synaptic labeling: *trans-Tango*, which induces both myrGFP and mtdTomato to be produced by certain presynaptic neurons and their post-synaptic partners through an engineered signaling pathway^77^. When *trans-Tango* was driven by *GAL4^SIFa.PT^*, we observed a significant overlap of 40-45% between SIFa neurons and SIFa trans-Tango signals in the PI and AG region (Fig. 7G and 7I; Fig. S7F and S7G). This suggests that SIFa neurons establish synaptic connections with each other, which supports the results obtained from the DenMark and syt.eGFP experiments (Fig. 2R).

SIFa neurons project to the protocerebral bridge (PB), which demonstrates several fundamental wiring principles within the central complex (CX), indicating that PB performs crucial computational tasks in the fly brain, such as the transformation of sensory (e.g., visual) input into locomotor commands^78^. SIFa neurons also project to noduli (No) in the CX, which are involved in olfactory processing and memory formation^79,80^, and the PRW region, where it is attached to the subesophageal ganglion (SOG), which contains the circuitry underlying feeding behavior and is involved in numerous other aspects of sensory processing and motor control^81^. Additionally, we discovered that SIFa neurons project to a small subset of OL neurons (Bottom panels in Fig. 7G). The most prominent projection of SIFa neurons is the AG of the VNC, which regulates abdominal muscles, the gastric system, and reproductive organs^82^. These findings imply that SIFa neurons project to an important brain region for extremely complicated computation and to a crucial VNC region for control of reproductive behavior, all of which modulate interval timing behavior based on the social context (Fig. 7L).

To examine the SIFa projections in further detail, we utilized *SIFaR^24F^*^06^*-GAL4*, which we recently identified as a key *GAL4* strain, which labels the essential VNC neurons expressing SIFaR for both LMD and SMD in AG^83^. The majority of *SIFaR^24F^*^06^ neurons in the brain project to the PI, SOG, and OL regions (GFP panels of Brain in Fig. 7J). In VNC, the majority of *SIFaR^24F^*^06^ neurons project to the AG region RFP signals J (Fig. 7J). We discovered that the projection of SIFa neurons in AG parallels the myrGFP signals of *SIFaR^24F^*^06^ neurons, indicating that these AG neurons are the direct synaptic target of SIFa neurons from the brain to the VNC to modulate interval timing. (Fig. 7K). Our results suggest that SIFa neurons in the PI of the brain and SIFaR neurons in the AG of the VNC are functionally linked for SIFa-SIFaR signaling (Fig. 7M).

## DISCUSSSION

In this study, we conducted a widely search to identify the neuropeptide SIFa that influences both male-specific interval timing behaviors in *D. melanogaster* (Table S1 and Document S1. Fig. S1–S5). We examined the association of SIFa-mediated signaling and its functional relevance to various behaviors including interval timing. The level of SIFa expression in four neurons adjacent to the PI region is essential for the generation of both LMD and SMD behaviors (Fig. 1 and S1). SIFa neurons extend their axons, rather than dendrites, throughout the entire brain and VNC, establishing robust synapses exclusively within the brain area (Fig. 2 and S2). SIFa produces several neurotransmitters, including glutamine, dopamine, and octopamine. However, only dopamine is essential for regulating SIFa-induced sleep, while all three neurotransmitters play a role in moderating SMD behavior and courtship activity (Fig. 3 and S3). The sNPF-sNPF-R peptidergic signaling provides input signals to SIFa neurons, which subsequently send back these signals to sNPF neurons through SIFaR. This feedback loop modifies the activity of LMD and SMD in particularly (Fig. 4 and S4). Of the various neuropeptidergic inputs to SIFa neurons, hunger and satiety signals from hugin and MIP do not affect LMD and SMD behaviors. However, Dsk inputs are shown to be crucial in regulating interval timing behaviors (Fig. 5 and S5). The elevated neuronal activity of SIFa neurons is linked to reduced mating duration as well as increased food intake, and vice versa. This suggests that the modulation of energy homeostasis-related behavioral change is dependent on the activity of SIFa neurons, as well as the co-transmission of octopamine and SIFa following this process (Fig. 6 and S6). The elevated neuronal activity of SIFa neurons is closely linked to reduced branching of SIFa axons at the postsynaptic level. Furthermore, the postsynaptic neurons of SIFa exhibit a significant degree of overlap in the CB, OL, and AG regions of the nervous system, where SIFaR is expressed (Fig. 7 and S7). Therefore, these findings highlight the crucial role of SIFa neuropeptides in regulating various behaviors in *D. melanogaster*, including interval timing, sleep, courtship, and feeding, through complex interactions with other neuropeptidergic systems.

Our study shows that SIFa neurons function as a central hub for integrating various internal signals and translating them into specific behavioral outputs, potentially offering novel insights into the regulation of complex behaviors in other organisms. In short, this study demonstrates that both input signals mediated by neuropeptides and output signals transmitted through SIFa-SIFaR communication or diverse neurotransmitters can modulate a range of complex behaviors related to energy homeostasis, including sleep, feeding, courtship, and interval timing. The SIFa neurons serve as a major hub for the integration of diverse inputs. In this study, we examined the fly SCope scRNA sequencing dataset to assess the expression of several neuropeptide receptors (NPRs) and neurotransmitter receptors (NTRs) in SIFa cells. To establish a control, we utilized gustatory receptors (GRs). SIFa neurons exhibit a high expression of NPRs and NTRs (Fig. S8A and S8B), indicating that they serve as a central hub for integrating different peptidergic and synaptic inputs in the brain.

While it has been reported that SIFa translates signals related to hunger and satiety through neuropeptides hugin and MIP^44^, we were unable to verify whether these neuropeptides affect interval timing behaviors through their receptor signaling. However, we discovered that sNPF functions as a crucial controller of interval timing behaviors through its receptor sNPF-R, which is expressed in SIFa neurons (as shown in Fig. 4 and S4). In addition to its involvement in interval timing behavior, sNPF also contributes to the regulation of sleep and feeding in *D. melanogaster*^57,58,61^. The activation of SIFa- and sNPF-expressing neurons using optogenetics has been observed to induce sleep in fruit flies, but the mechanisms involved are quite distinct^59,84^. However, the exact functions of these neurons in regulating sleep are not yet completely understood. We demonstrated the cooperative interaction between the SIFa and sNPF systems through the establishment of a feedback loop, facilitated by the relay of neuropeptides and their subsequent binding to their respective receptors in specific neural circuits. Our studies revealed that SIFa neurons, situated in the brain, transmit signals related to internal states to sNPF neurons that express SIFaR in the VNC. This conclusion is substantiated by our discovery that SIFa establishes robust synaptic connections with sNPF-R neurons in the LAL, a brain region that crucially regulates behavior by linking the brain and thoracic motor center with a group of descending neurons. Further investigations into the downstream targets and modulatory processes of SIFa signaling offer promise for explaining its full contribution to behavioral regulation.

## MATERIALS AND METHOD

### Fly Stocks and Husbandry

*Drosophila Melanogaster* were raised on cornmeal-yeast medium at similar densities to yield adults with similar body sizes. Flies were kept in 12h light: 12h dark cycles (LD) at 25°C (ZT 0 is the beginning of the light phase, ZT12 beginning of the dark phase) except for some experimental manipulation (experiments with the flies carrying tub-GAL80^ts^). Wild-type flies were Canton-S(CS).

Following lines used in this study, *Canton-S* (#64349)*, w1118* (#3605)*, SIFa^2A-GAL4^* (#84690)*, ple-RNAi* (#25796)*, Trhn-RNAi* (#25842*), ChAT-RNAi* (#25856)*, Gad1-RNAi* (#28079)*, VGlut-RNAi* (#27538)*, hdc-RNAi* (#30489)*, SIFa-RNAi* (#60484)*, sNPF-RNAi* (#25867)*, sNPF-R-RNAi* (#27507)*, SIFaR-RNAi* (#34947)*, SIFaR^24F^*^06^*-GAL4* (#49087)*, sNPF-R^64B^*^11^*-GAL4* (#41288)*, sNPF-R^64H^*^09^*-GAL4* (#46547)*, CCKLR17D1-RNAi* (#27494)*, CCKLR17D3-RNAi* (#28333)*, Dimm-RANi* (#38530)*, UAS>stop>mCD8GFP* (#30125)*, fru^FLP^* (#66870)*, UAS-KCNJ2* (#6596)*, UAS-TNT* (#28838)*, UAS-NaChBac-EGFP* (#9466)*, SIFa[1]* (#80696)*, SIFa[2]* (#80697)*, UAS-myr-GFP* (#32197)*, UAS-CD4tdGFP* (#35839)*, UAS-Redstinger* (#32222)*, UAS-mCD8RFP, lexAop-mCD8GFP* (#32229), *lexAop-nSyb-spGFP1-10, UAS-CD4-spGFP11* (#64315), *UAS-mCD8-RFP* (#61679), *UAS-myrGFP.QUAS-mtdTomato-3xHA, trans-Tango* (#77124)*, elav^c^*^155^*; UAS-Dicer* (#25750), *UAS-Denmark, UAS-syt.eGFP* (#33065), *UAS-mCD8RFP, lexAop-mCD8GFP* (#32229) were obtained from the Bloomington *Drosophila* Stock Center at Indiana University. The following lines, *CrzR-RNAi* (#108506), *PK2-R2-RNAi* (#44871), *CCHa1-R-RNAi* (#103055). *InR-RNAi* (#991), *UAS-TrpA1* (#61504), *UAS-ShiBirets* (#66600), *UAS-dsx^F^*(#44223), *UAS-tra^F^*(#4590), *UAS-sxl* (#55769) were obtained from the Vienna Drosophila Resource Center. The following lines, *SIFa^2A-lexA^* (#FBA00106), *SIFaR^2A^-GAL4* (#FBA00102)*, UAS-t-SIFa-SEC, UAS-t-SIFa-ML, SIFa^PT^-lexA* (see methods: Generation of transgenic flies and SIFa t-Peptide (t-SIFa) generation). The GAL4*^SIFa.PT^* was a gift from Jan A. Veenstra. *UAS-OrkΔC* were obtained from Nitabach MN.

To reduce the variation from genetic background, all flies were backcrossed for at least 3 generations to CS strain. All mutants and transgenic lines used here have been described previously.

### Mating duration assay

The mating duration assay in this study has been reported^34,71,85^. To enhance the efficiency of the mating duration assay, we utilized the *Df (1)Exel6234* (DF here after) genetic modified fly line in this study, which harbors a deletion of a specific genomic region that includes the sex peptide receptor (SPR)^86,87^. Previous studies have demonstrated that virgin females of this line exhibit increased receptivity to males^87^. We conducted a comparative analysis between the virgin females of this line and the CS virgin females and found that both groups induced SMD. Consequently, we have elected to employ virgin females from this modified line in all subsequent studies. For naïve males, 40 males from the same strain were placed into a vial with food for 5 days. For single reared males, males of the same strain were collected individually and placed into vials with food for 5 days. For experienced males, 40 males from the same strain were placed into a vial with food for 4 days then 80 DF virgin females were introduced into vials for last 1 day before assay. 40 DF virgin females were collected from bottles and placed into a vial for 5 days. These females provide both sexually experienced partners and mating partners for mating duration assays. At the fifth day after eclosion, males of the appropriate strain and DF virgin females were mildly anaesthetized by CO_2_. After placing a single female in to the mating chamber, we inserted a transparent film then placed a single male to the other side of the film in each chamber. After allowing for 1 h of recovery in the mating chamber in 25℃ incubators, we removed the transparent film and recorded the mating activities. Only those males that succeeded to mate within 1 h were included for analyses. Initiation and completion of copulation were recorded with an accuracy of 10 sec, and total mating duration was calculated for each couple. All assays were performed from noon to 4pm. We conducted blinded studies for every test^98,99^.

### Quantitative analysis of fluorescence intensity

To quantify the calcium level and synaptic intensity in microscopic images, we introduced ImageJ software^88^. We initially employed ImageJ’s ‘Measure’ feature to calculate average pixel intensity across the entire image or in user-specified sections, and the ‘Plot Profile’ feature to create intensity profiles across areas. To maximize precision, we converted color images to grayscale before analysis. Thresholding methods were also utilized to produce binary images that accurately outlined areas of interest, with pixel intensities of 255 (white) assigned to regions of interest and 0 (black) to the background. Intensity values from the binary image were then transferred to the corresponding locations in the original grayscale image to obtain precise intensity measurements for each object. The ‘Display Results’ feature provided comprehensive data for each object, including average intensity, size, and other relevant statistics. To normalize for fluorescence differences between ROIs, GFP fluorescence for CaLexA and GRASP was normalized to nc82. All specimens were imaged under identical conditions.

### Generation of transgenic flies

To generate the *SIFa^PT^-lexA* driver, the putative promotor sequence of the gene was amplified by PCR using wild-type genomic DNA as a template with the following primers GCCAATTGGCTGAATCTCCTGACCCTCA and GCAGATCTCTTGCAGTTTTCGGTGAGC as mentioned before^89^. The amplified DNA fragment (1482 base pairs located immediately upstream of the *SIFa* coding sequence) was inserted into the E2 Enhancer-lexA vector. This vector, supplied by Qidong Fungene Biotechnology Co., Ltd. (http://www.fungene.tech/), is a derivative of the pBPLexA::p65Uw vector (available at https://www.addgene.org/26231). The insertion was achieved by digesting the fragment and the vector with EcoRI and XbaI restriction enzymes to create compatible sticky ends. The genetic construct was inserted into the attp2 site on chromosome III to generate transgenic flies using established techniques, a service conducted by Qidong Fungene Biotechnology Co., Ltd.

### SIFa t-Peptide (t-SIFa) generation

To generate the *UAS-t-SIFa-ML* and *UAS-t-SIFa-SEC* driver in Fig. 1H and 1K, t-peptide cDNAs were chemically synthesized with optimal *Drosophila* codon usage and with an optimal *Drosophila* Kozak translation initiation site upstream of the start methionine (CAAA)^90^. Encoded t-peptide is as follows: *t-SIFa-ML*, MSALLILALVGAAVAAYRKPPFNGSIFGNEQKLISEEDLGNGAGFATPVTLAL VPALLATFWSLL. These cDNAs were cloned into the pUAS-attB vector; For generation of transgenic *Drosophila*, Vectors was injected into the embryos of flies. The genetic construct was inserted into the attp40 site on chromosome II to generate transgenic flies using established techniques, a service conducted by Qidong Fungene Biotechnology Co., Ltd. (http://www.fungene.tech/).

### Antibody preparation

To generate the SIFa-antibody in Fig. S1B, the peptide sequence RDKKTLKIKMKLFI of SIFa was synthesized and conjugated to Keyhole Limpet Hemocyanin (KLH). The efficiency of the conjugation was confirmed by SDS-PAGE analysis. Healthy rabbits were immunized via multiple subcutaneous injections with the peptide-KLH conjugate mixed with Freund’s adjuvant. The immunization schedule included a primary immunization and two booster injections, each separated by three weeks. Two weeks after the final booster, serum was collected from the rabbits via cardiac puncture. The serum was purified using affinity chromatography, and the antibody titer and specificity were determined by ELISA. Antibody synthesis was performed by Wuhan Vapol Bioscience Company (http://www.vapolbio.com).

### Courtship Assay

Courtship assay was performed as previously described^91^, under normal light conditions in circular courtship arenas 11 mm in diameter, from noon to 4 p.m. Courtship latency is the time between female introduction and the first obvious male courtship behavior, such as orientation coupled with wing extensions. Once courtship began, courtship index was calculated as the fraction of time a male spent in any courtship-related activity during a 10 min period or until mating occurred.

### Immunostaining

The dissection and immunostaining protocols for the experiments are described elsewhere. After 5 days of eclosion, the *Drosophila* brain will be taken from adult flies and fixed in 4% formaldehyde at room temperature for 30 minutes. The sample will be washed three times (5 minutes each) in 1% PBT and then blocked in 5% normal goat serum for 30 minutes. The sample will next be incubated overnight at 4°C with primary antibodies in 1% PBT, followed by the addition of fluorophore-conjugated secondary antibodies for one hour at room temperature. The brain will be mounted on plates with an antifade mounting solution (Invitrogen) for imaging purposes.

Samples were imaged with Zeiss LSM880. Antibodies were used at the following dilutions: Chicken anti-GFP (1:500, Invitrogen), mouse anti-nc82 (1:50, DSHB), rabbit anti-DsRed (1:500, Rockland Immunochemicals), Alexa-488 donkey anti-chicken (1:200, Jackson ImmunoResearch), Alexa-555 goat anti-rabbit (1:200, Invitrogen), Alexa-647 goat anti-mouse (1:200, Jackson ImmunoResearch), Rat anti-elav (1:200, DSHB), Mouse anti-repo (1:200, DSHB), Rabbit anti-SIFa (1:200, see methods: Antidody preparation).

### RNA extraction and cDNA synthesis

RNA was extracted from 50 preparations of 5-day-old CS males using the RNA isolation kit (Vazyme), following the manufacturer’s protocol. And first-strand cDNA was synthesized from 1 μg of RNA template with random primers using SPARK script Ⅱ RT plus kit (SparkJade).

### Quantitative RT-PCR

The expression levels of *SIFa* in naïve, exp and single conditions were analyzed by quantitative real-time RT-PCR with SYBR Green qPCR MasterMix kit (Selleckchem). The primers of RT-PCR are F:5’-aagcaggagagcgagttcag-3’; R: 5’-ttcgccttgttttgtcacag-3’^92^. qPCR reactions were performed in triplicate, and the specificity of each reaction was evaluated by dissociation curve analysis. Each experiment was replicated three times. PCR results were recorded as threshold cycle numbers (Ct). The fold change in the target gene expression, normalized to the expression of internal control gene (GAPDH) and relative to the expression at time point 0, was calculated using the 2 ^−ΔΔCT^ method as previously described^93^. The results are presented as the mean ± SD of three independent experiments.

### Colocalization analysis

To perform colocalization analysis of multi-color fluorescence microscopy images in this study, we employed ImageJ software. Initially, merge the individual channels of the fluorescence image to create a composite view. Following this, access the ‘Image’ menu, select ‘Type’, and then choose ‘RGB Color’ to ensure the image is displayed in the correct color format. Next, navigate to ‘Image’, then ‘Adjust’, and select ‘Color Threshold’. Adjust the threshold settings to isolate the yellowish pixels indicative of colocalization, and confirm the selection by clicking the ‘Select’ button.

Subsequently, proceed to ‘Analyze’, followed by ‘Measure’. This action will open a new window displaying the measurement data for the selected area. Record the ‘Area’ value provided in the measurement window, as this corresponds to the area of colocalization between the two fluorophores. To calculate the percentage of the colocalized area relative to the total area of the fluorophore of interest (e.g., GFP or RFP), readjust the threshold settings to encompass the entire fluorophore area and click ‘Select’ again. Afterward, repeat the measurement process by going to ‘Analyze’ and then ‘Measure’. This will yield the total area value for the respective fluorophore. Finally, to determine the colocalization percentage, divide the area value of the colocalized region by the total area value of the GFP or RFP region. This calculation provides the colocalization efficiency within the specified region.

Additionally, we displayed images of fluorescence and overlapping areas in the male fly brain and/or VNC, processed with ImageJ software using a threshold function to differentiate fluorescence from background. All specimens were imaged under identical conditions.

### Locomotion assay

To detect and quantify the activity of flies, we have developed the Fly Trajectory Dynamics Tracking (FlyTrDT) software. This is an open-source, custom-written Python program that utilizes the free OpenCV machine vision library and the Python Qt library. The FlyTrDT software simultaneously records the trajectory information of each fly and calculates various indicators of the group at a certain period. For each frame acquired, the moving fly is segmented using the binarization function from the OpenCV library. Subsequently, a Gaussian blur and morphological closing and opening operations were performed on the extracted foreground pixels to consolidate detected features and reducing false positives and negatives. Finally, the extraction of fly outlines was achieved using the contour detection algorithm in the OpenCV library.

### Sleep assay of single fly

Adult male flies (aged 3-7 days) were loaded into wells of white 96-well Microfluor 2 plates (Fishier) containing 300ul food (5% sucrose and 1% agar). The fly boxes were placed in an incubator to control temperature precisely. A webcam tracked the locomotion of flies at 10s intervals, and the data were used for sleep analysis^94^. All experiments were replicated at least three times.

### Sleep assay of group reared flies

To investigate *Drosophila* circadian rhythms, we utilized the DAM system^95^, which records infrared beam breaks as flies move through glass tubes. Each DAM monitor tracked 32 individual flies, with experiments scaling to hundreds. Flies were housed in tubes with food for extended monitoring, and drugs were added to modulate behavior. Monitors were placed in incubators controlling temperature, humidity, and light to study environmental responses and transgene expression. We developed ShinyR-DAM, a free, open-source, cloud-based application for rapid data analysis, producing customizable plots and CSV files. ShinyR-DAM streamlines the analysis of LD and DD conditions, eliminating complex installations and manual dead fly detection. It leverages the R programming language’s Shiny framework, making it accessible and customizable for researchers.

### Blue dye assay

For colorimetric food intake assay, flies were starved in PBS-containing vials for 2h and fed for 15min in vials containing 0.05% FD&C Blue dye, 7% sucrose and 5% yeast. The flies were frozen, homogenized in PBS, and centrifuged twice for 25min each. The supernatant was measured at 625nm. Each experiment consisted of 20 flies and the assay was repeated three times^96^.

### Ex-Q assay

2 flies were placed in a tube with 5% sucrose, 5% yeast and 1% agar with 0.5% erioglaucine disodium blue dye. After 24 h, flies were homogenized in tubes with its excreta and centrifuged for 15 min in 13,000 rpm. The supernatant was measured at 630 nm. For TrpA1 and shi^ts^ thermogenetic experiments, flies were maintained and starved in 20°C and transferred to 30°C 30 mins before testing.

### Single-nucleus RNA-sequencing analyses

The snRNAseq dataset analyzed in this paper is published^97^ and available at the Nextflow pipelines (VSN, https://github.com/vib-singlecell-nf), the availability of raw and processed datasets for users to explore, and the development of a crowd-annotation platform with voting, comments, and references through SCope (https://flycellatlas.org/scope), linked to an online analysis platform in ASAP (https://asap.epfl.ch/fca).

### Gene expression analysis

Single-cell RNA sequencing (scRNA-seq) data from the *Drosophila melanogaster* were obtained from the Fly Cell Atlas website (https://doi.org/10.1126/science.abk2432). Our analysis employed UMI (Unique Molecular Identifier) data extracted from the 10x VSN Head (Stringent) loom file, encompassing a total of 100,527 cells. The Seurat (v4.2.2) package (https://doi.org/10.1016/j.cell.2021.04.048) was utilized for data analysis. Genes with a Log2 fold change surpassing 0 were retained.

### Statistical Tests

Statistical analysis of mating duration assay was described previously^98–100^. More than 50 males (naïve, experienced and single) were used for mating duration assay. Our experience suggests that the relative mating duration differences between naïve and experienced condition and singly reared are always consistent; however, both absolute values and the magnitude of the difference in each strain can vary. So, we always include internal controls for each treatment as suggested by previous studies. Therefore, statistical comparisons were made between groups that were naïvely reared, sexually experienced and singly reared within each experiment. As mating duration of males showed normal distribution (Kolmogorov-Smirnov tests, p > 0.05), we used two-sided Student’s t tests. The mean ± standard error (s.e.m) (**** = p < 0.0001, *** = p < 0.001, ** = p < 0.01, * = p < 0.05). All analysis was done in GraphPad (Prism). Individual tests and significance are detailed in Fig. legends. Besides traditional t-test for statistical analysis, we added estimation statistics for all MD assays and two group comparing graphs. In short, ‘estimation statistics’ is a simple framework that—while avoiding the pitfalls of significance testing—uses familiar statistical concepts: means, mean differences, and error bars. More importantly, it focuses on the effect size of one’s experiment/intervention, as opposed to significance testing. In comparison to typical NHST plots, estimation graphics have the following five significant advantages such as (1) avoid false dichotomy, (2) display all observed values (3) visualize estimate precision (4) show mean difference distribution. And most importantly (5) by focusing attention on an effect size, the difference diagram encourages quantitative reasoning about the system under study. Thus, we conducted a reanalysis of all of our two group data sets using both standard t tests and estimate statistics. In 2019, the Society for Neuroscience journal eNeuro instituted a policy recommending the use of estimation graphics as the preferred method for data presentation.

## ACKNOWLEDGMENTS

We thank Dr. Jan A. Veenstra (University of Bordeaux) for sharing SIFa^PT^-GAL4 driver, Drs. Yuh Nung Jan and Lily Yeh Jan (UCSF, USA) for helpful comments and support on this paper. We are also very appreciative to the colleagues who supplied us with several fly strains: Dr. Wei Zhang (Tsinghua University), Fang Guo (Zhejiang University), and Dr. Yufeng Pan (Southeast University) and Drs. Young-Joon Kim and Sung-Eun Yoon (Korea Drosophila Resource Center, KDRC). The authors would like to express their gratitude to NVIDIA Academic Hardware Grant Program for providing GPU for behavioral analysis. This research was supported a University of Ottawa Startup grant 602496 to WJK, Startup funds from HIT Center for Life Science to WJK, a University of Ottawa Interdisciplinary Research Group Funding Opportunity (IRGFO stream 1 and 2) grants 148101 and 148747 to WJK, a Natural Sciences and Engineering Research Council of Canada (NSERC) Discovery grant (reference: 211406) to WJK, a University of Ottawa Brain and Mind Research Institute/Center for Neural Dynamics Open call project grant 150950 to WJK, a Mitacs Globalink Research Internship Program grant 17268 to WJK. This research was also supported by the Brain Pool Program of the National Research Foundation in Korea grant ZYM5041911 to WJK, Burroughs Wellcome Fund Collaborative Research Travel Grants (reference: 1017486) to WJK and a NVIDIA Academic Hardware Grant Program to WJK. The funders had no role in study design, data collection and analysis, decision to publish, or preparation of the manuscript. SGL received salary from the ‘University of Ottawa Startup grant to WJK’ and HM from the ‘Startup funds from HIT Center for Life Science to WJK’.

## CONTRIBUTIONS

**Conceptualization:** Woo Jae Kim.

**Data curation:** Yutong Song, Tianmu Zhang, Tae Hoon Ryu, Zekun Wu, Kyle Wong, Justine Schweizer, Woo Jae Kim.

**Formal analysis:** Yutong Song, Tianmu Zhang, Tae Hoon Ryu, Wenjing Li, Yanying Sun, Xiaoli Zhang, Zekun Wu, Kyle Wong, Justine Schweizer, Khoi-Nguyen Ha Nguyen, Alex Kwan, and Woo Jae Kim.

**Funding acquisition:** Woo Jae Kim.

**Investigation:** Woo Jae Kim.

**Methodology:** Tae Hoon Ryu, Woo Jae Kim.

**Project administration:** Woo Jae Kim.

**Resources:** Woo Jae Kim.

**Supervision:** Woo Jae Kim.

**Validation:** Yutong Song, Tianmu Zhang, Woo Jae Kim.

**Visualization:** Zekun Wu, Woo Jae Kim.

**Writing – original draft:** Woo Jae Kim.

**Writing – review & editing:** Yutong Song, Kweon Yu, Woo Jae Kim.

## DECLARATION OF GENERATIVE AI AND AI-ASSISTED TECHNOLOGIES IN THE WRITING PROCESS

During the creation of this work, the author(s) utilized QuillBot to rephrase English sentences, verify English grammar, and detect plagiarism, as none of the authors of this paper are native English speakers. After using this tool/service, the author(s) reviewed and edited the content as needed and take(s) full responsibility for the content of the publication.

---

Supplementary information is available at cell discovery’s website.

**Movie 1.** 3D immunofluorescent Visualization of SIFa Neurons in the *Drosophila* Brain.

Flies expressing *GAL4^SIFa.PT^* together with *UAS-RedStinger* were immunostained with anti-elav (green), anti-repo (blue) and anti-SIFa(red) antibodies.

**Table S1.** Summary of neuropeptide receptors *RNAi* screening with *elav^c^*^155^ driver

**Fig. S1.**
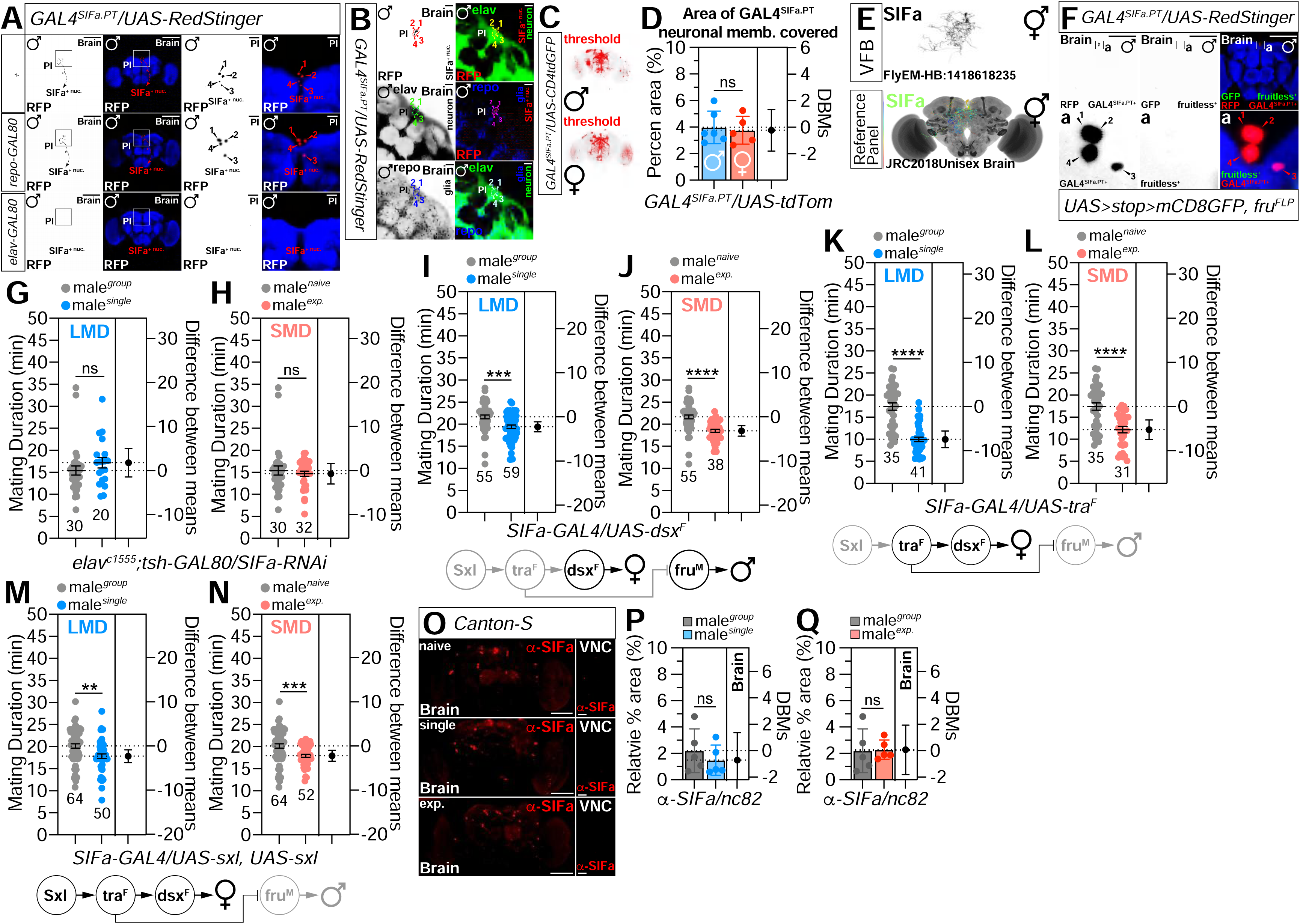
*SIFa*-expressing neurons are not linked to sexual dimorphism. (A) Flies expressing *GAL4^SIFa.PT^ /+*(upper) or *repo-GAL80; GAL4^SIFa.PT^* (middle) or *elav-GAL80; GAL4^SIFa.PT^* (bottom) together with *UAS-RedStinger* were immunostained with anti-DsRed (red), and nc82 (blue) antibodies. Scale bars represent 100 mm in the left two panels and 10 mm in the magnified right panels. (B) Flies expressing *GAL4^SIFa.PT^* together with *UAS-RedStinger* were immunostained with anti-elav (green), anti-repo (blue) and anti-SIFa(red) antibodies. Arrowheads: cell body locations. Scale bars represent 10 µm. (C) The threshold of GFP fluorescence in male and female fly brain was marked by threshold function of ImageJ. (D) Quantification of male and female brain regions covered by the *SIFa* cell membrane. (E) Expression pattern of *SIFa* in Virtual Fly Brain (VFB). (F) Flies expressing *GAL4^SIFa.PT^*drivers together with *UAS>stop>mCD8GFP; fru^FLP^* was immunostained with anti-GFP (green), anti-RFP (red) and nc82 (blue) antibodies. Scale bars represent 100 µm. (G-H) MD assays for *elav-GAL4* drivers mediated knockdown of *SIFa* in tsh-negative neurons via *tsh-GAL80; SIFa-RNAi* (I-N) MD assays for *GAL4^SIFa.PT^* drivers mediated expression of female form of doublesex (*UAS-dsx^F^*), transformer (*UAS-tra^F^*), or sex lethal (*UAS-sxl*). (O) The brains of CS *Drosophila* (left panel) and the VNC (right panel) were subjected to immunostained with anti-SIFa antibodies (red channel) under naïve, single, and exp conditions. Scale bars represent 100 µm in brain and 10 µm . (P-Q) Quantification of *SIFa* signals in Fig. S1O.

**Fig. S2.**
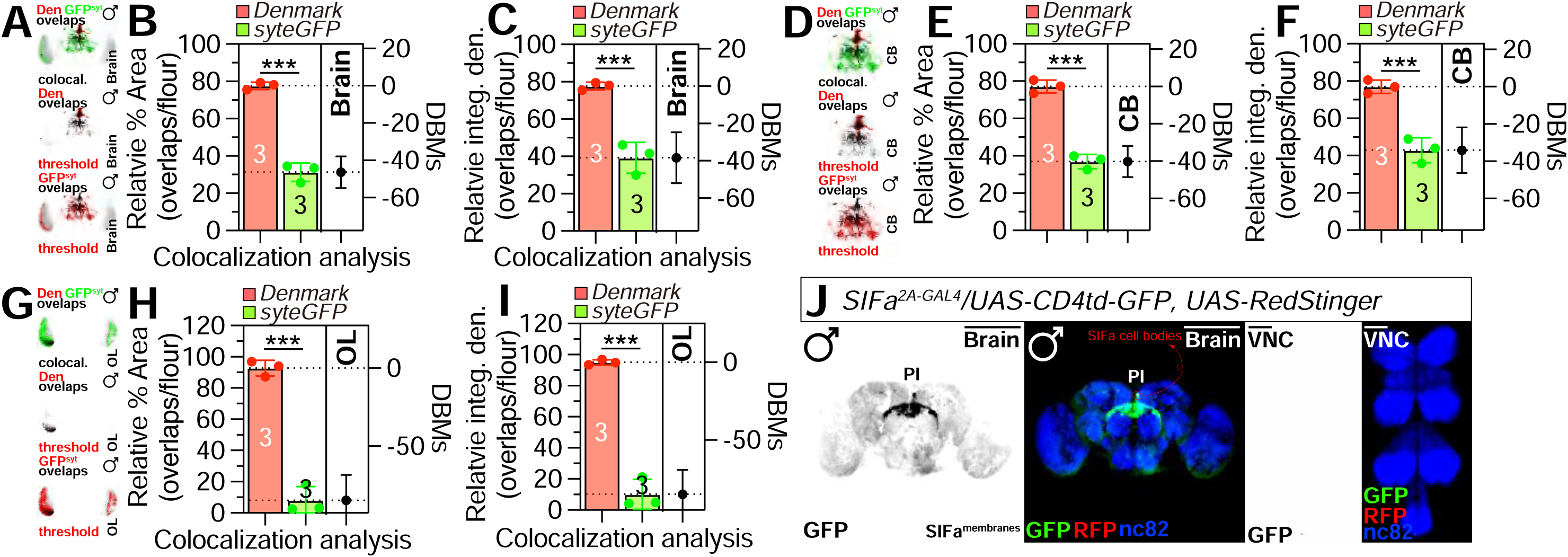
Presynaptic terminals are arborized more than dendrites of *SIFa* neurons. (A-I) Colocalization analysis of dendritic and presynaptic terminals of neurons labeled via *GAL4^SIFa.PT^* in the brain(A-C), CB(D-F) and OL(G-I). See the MATERIALS AND METHODS for a detailed description of the colocalization analysis used in this study. (J) Male flies expressing *SIFa^2A-GAL4^* together with *UAS-CD4tdGFP* and *UAS-RedStinger* were immunostained with anti-GFP (green), anti-RFP (red), nc82 (blue) antibodies. Scale bars represent 100 µm.

**Fig. S3.**
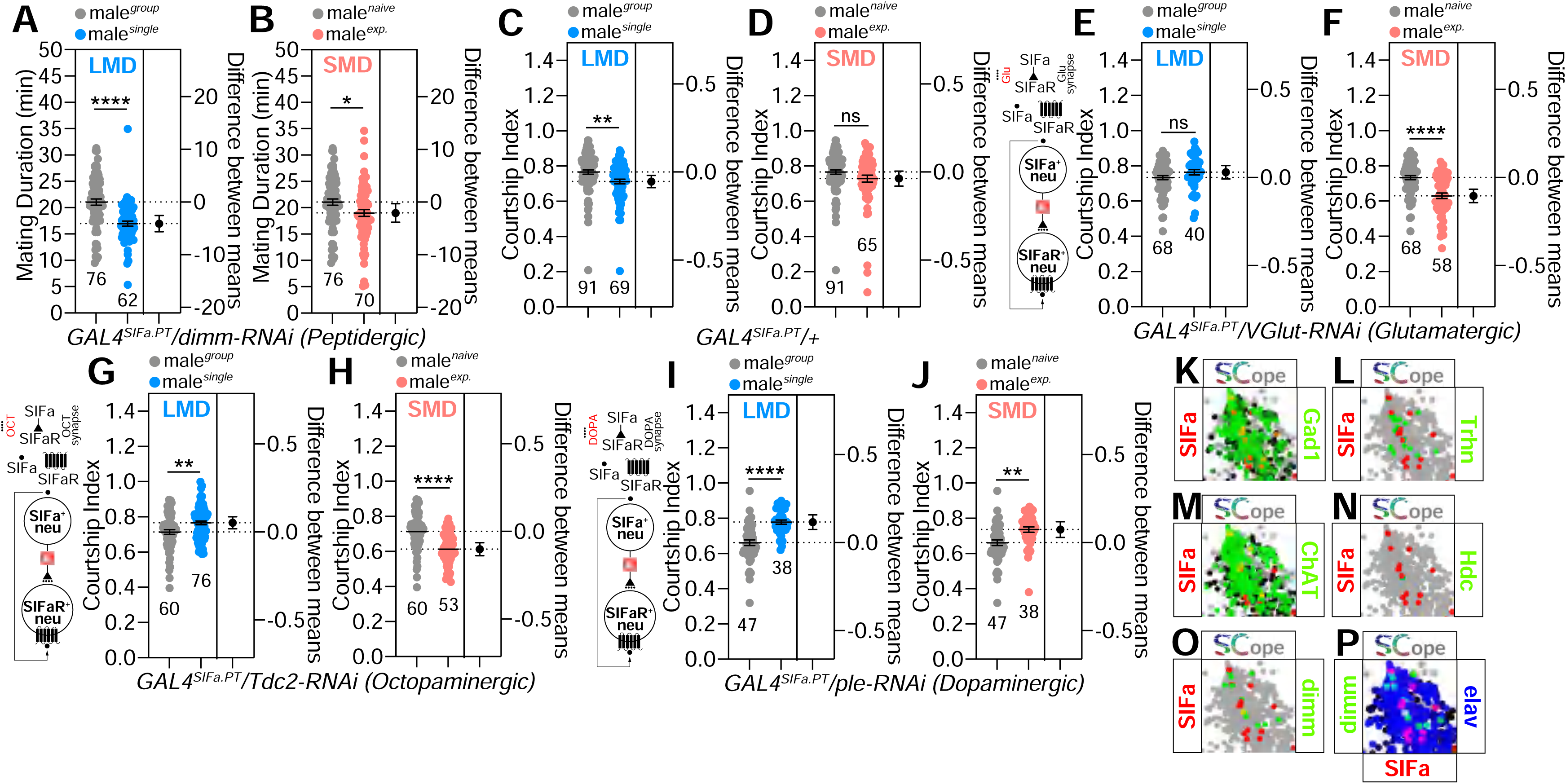
Sexual experience didn’t reduce male courtship behavior. (A-B) MD assays for *GAL4^SIFa.PT^* mediated knockdown of *dimm* (peptidergic) *via dimm-RNAi.* (C-D) courtship index of control flies in group, single and exp conditions. (E-J) courtship index of flies for *GAL4* mediated knockdown of *VGlut* (Glutamatergic), *Tdc2* (Octopaminergic) and *Ple* (Dopaminergic) via *VGlut-RNAi*, *Tdc2-RNAi* and *ple-RNAi* using the *GAL4^SIFa.PT^*. (K-O) Fly SCope single-cell RNA sequencing data of cells co-expressing *SIFa* together with *Gad1*, *Trhn*, *ChaT*, *Hdc*, *dimm* or with *dimm* and *elav*.

**Fig. S4.**
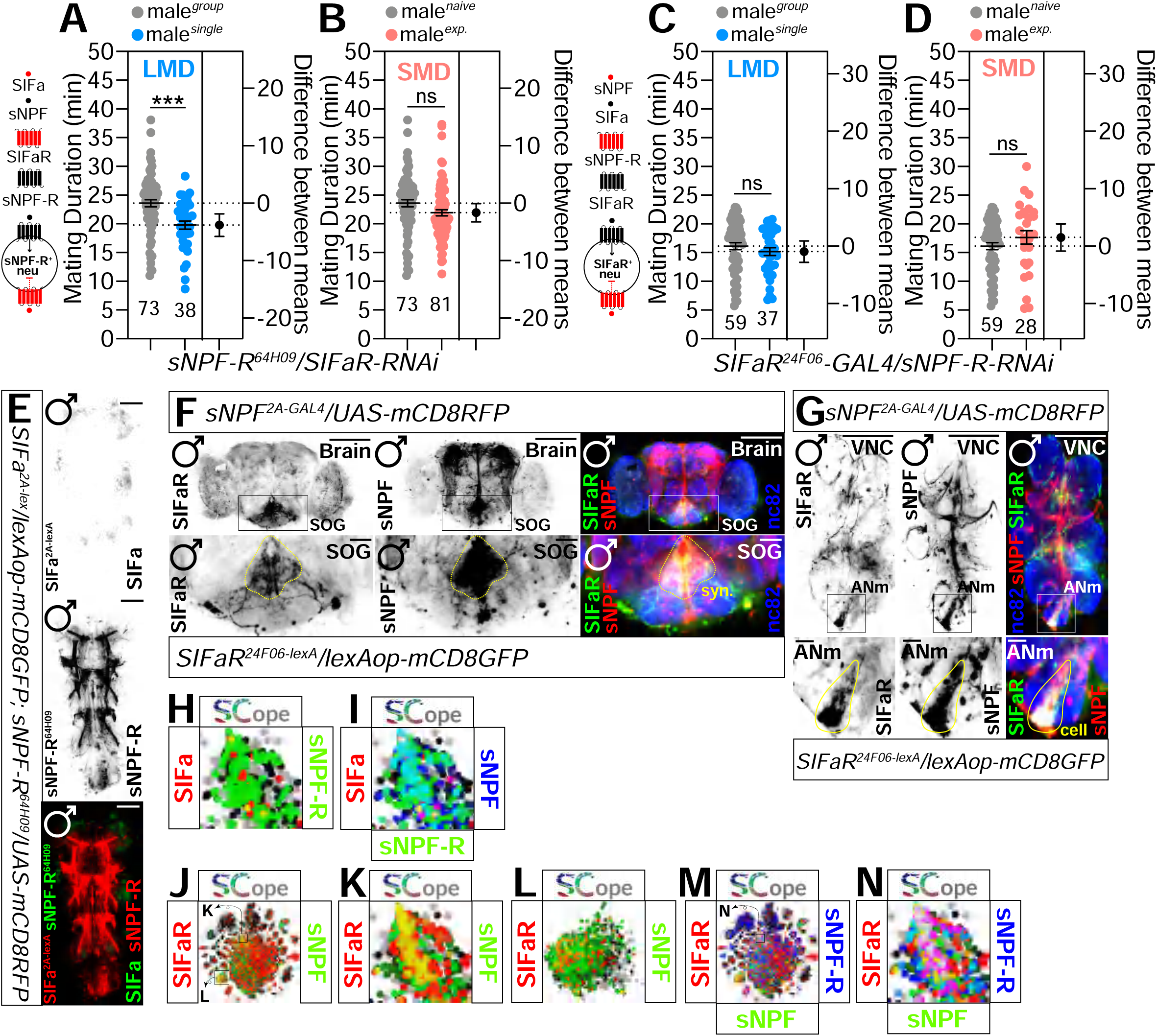
*SIFa* and *sNPF* collaborate to regulate interval timing behaviors. (A-D) MD assays for *GAL4* mediated knockdown of *SIFaR* and *sNPF-R* via *SIFaR-RNAi* and *sNPF-R-RNAi* using the *sNPF-R^64H09^-GAL4* driver (A-B), and *SIFaR^24F06^-GAL4* driver (C-D). (E) Male VNC of flies expressing *sNPF-R^64H09^-GAL4* and *SIFa^T2A-lexA^* drivers together with *UAS-mCD8RFP* and *lexAop-mCD8GFP* was immunostained with anti-GFP (green), anti-DsRed (red) and anti-nc82 (blue) antibodies. The top two panels are presented as a grey scale to clearly show the membrane expression patterns of *SIFa+* neurons in the adult labeled by *sNPF-R^64H09^-GAL4* driver. Scale bars represent 100 μm. (F-G) Male brain and VNC of flies expressing *sNPF^2A-GAL4^* together with *UAS-mCD8RFP* was immunostained with anti-GFP (green), anti-DsRed (red) and anti-nc82 (blue) antibodies. Scale bars represent 100 μm. areas outlined by black boxes are enlarged in the bottom panel. Scale bars represent 50 μm. (H-N) Fly SCope single-cell RNA sequencing data of cells co-expressing *SIFa* together with *sNPF-R*(H) *sNPF* and *sNPF-R*(I), *SIFaR* together with *sNPF* (J-K), and with *sNPF* and *sNPF-R*(M-N).

**Fig. S5.**
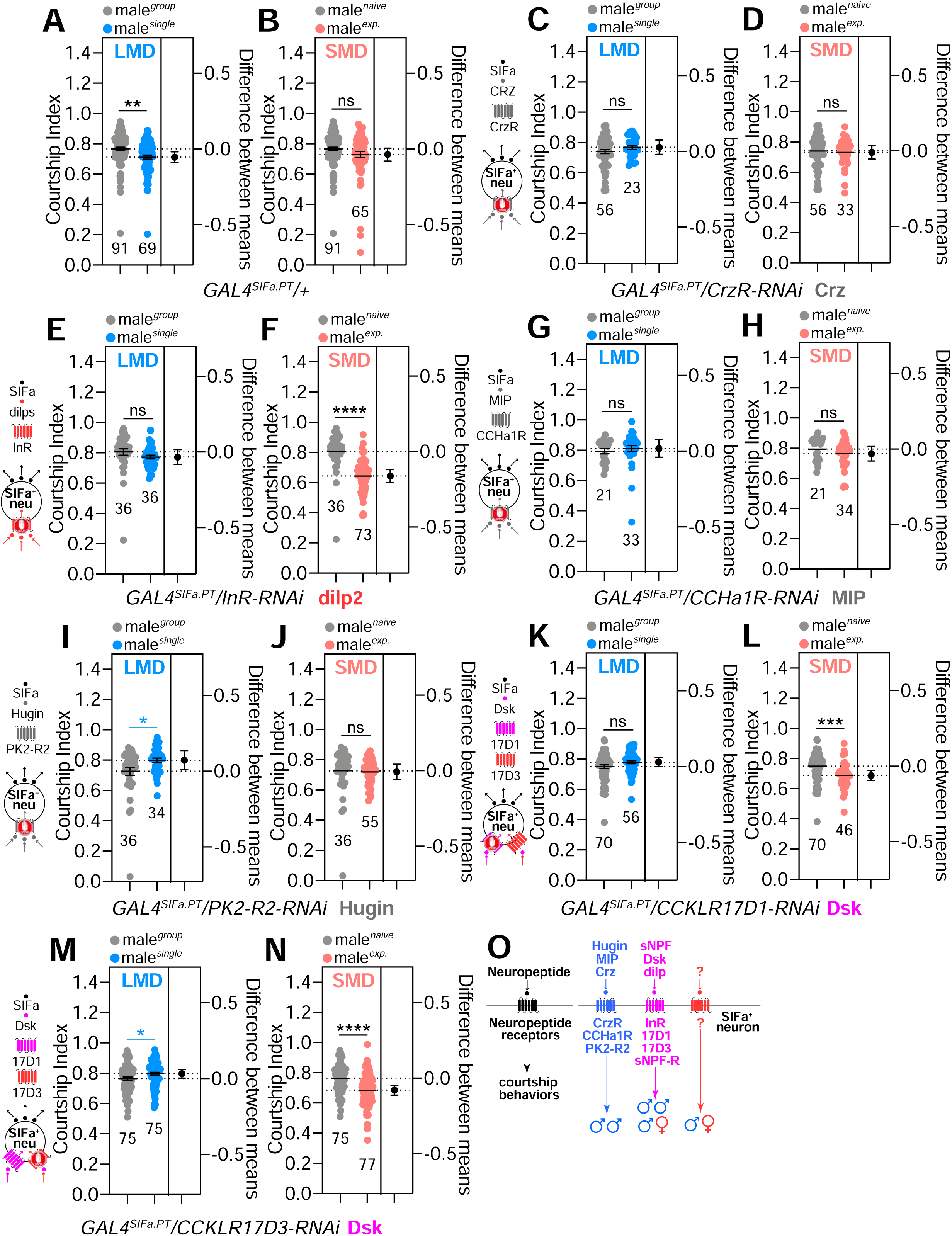
Various peptidergic inputs to *SIFa* influence courtship behavior. (A-B) courtship index of control flies in group, single and exp conditions. (C-N) courtship index of flies for *GAL4* mediated knockdown of *CrzR*, *InR*, *CCHa1R*, *PK2-R2*, *CCKLR17D1* and *CCKLR17D3* via *CrzR-RNAi*, *InR-RNAi*, *CCHa1R-RNAi*, *PK2-R2-RNAi*, *CCKLR17D1-RNAi* and *CCKLR17D3-RNAi* using the *GAL4^SIFa.PT^*. The diagram of the relationship between *SIFa* neurons and the neuropeptide receptors is on the left. (O) Schematic representation of the regulatory mechanisms by which neuropeptides modulate courtship behavior via their corresponding receptors.

**Fig. S6.**
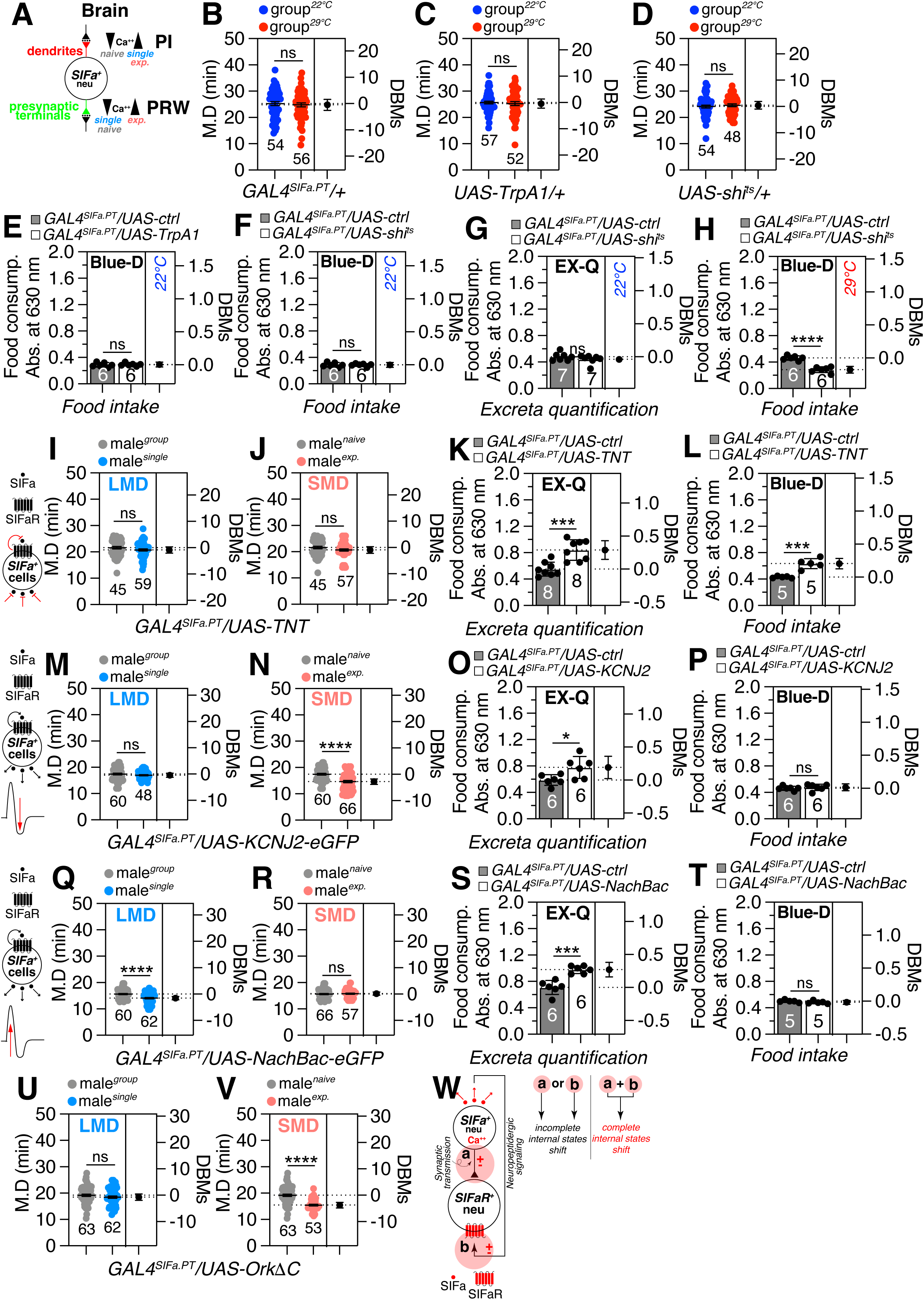
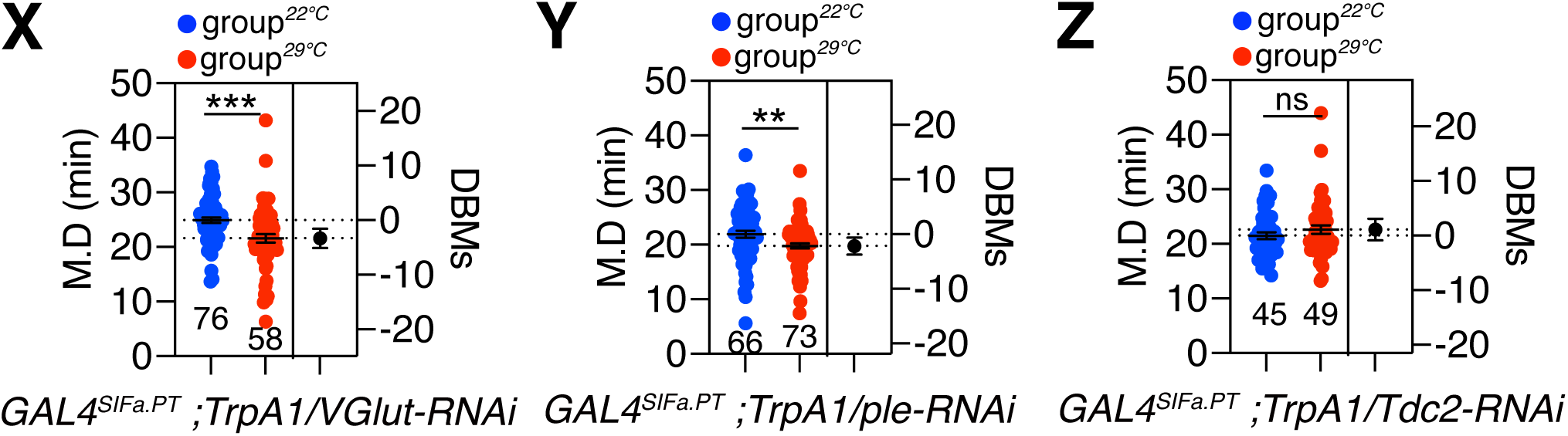
Deactivating *SIFa* neurons may reduce food intake. (A) Alterations in Ca2+ concentrations within *SIFa* neurons in response to social isolation and sexual experience exhibit a comparable yet distinct profile. (B-D) Genetic control of MD assay of flies expressing *TrpA1*(C) and shi^ts^ (D) in *SIFa* neurons at the activation and deactivation temperature. (E) Blue-Dye assay of flies expressing *GAL4^SIFa.PT^* driver together with *UAS-TrpA1* in 22℃. (F-H) Blue-Dye and EX-Q assay of flies expressing *GAL4^SIFa.PT^* driver together with *UAS-shi^ts^* in 22℃ and 29℃. (I-J) MD assays for *GAL4^SIFa.PT^* meditated the inactivation of synaptic transmission of *SIFa* neurons using *UAS-TNT*. (M-N, Q-R, U-V) Mating duration assays of *UAS-KCNJ2-eGFP*(K-L)*, UAS-NachBac-eGFP*(O-P) and *UAS-OrkΔC* crossed with *GAL4^SIFa.PT^*. (K-L, O-P, S-T) 24-h excreta quantification of males expressing *GAL4^SIFa.PT^*driver together with *UAS-TNT* (inactivation of synaptic transmission), *UAS-KCNJ*(M) and *UAS-NachBac* (Q) on yeast-sugar medium. Blue-Dye assay of male flies expressing *GAL4^SIFa.PT^* driver together with *UAS-TNT* (inactivation of synaptic transmission, J), *UAS-KCNJ* (N) and *UAS-NachBac* (R). (W) Schematic representation of the regulatory interactions between *SIFa* and *SIFaR*, in the context of modulating internal states. (X-Z) Analysis of mating duration in male *Drosophila* expressing *GAL4^SIFa.PT^* drivers, in conjunction with the heat-sensitive *Drosophila* cation channel *TrpA1* and *VGlut-RNAi*, *ple-RNAi* and *Tdc2-RNAi*, following temperature shifts.

**Fig. S7.**
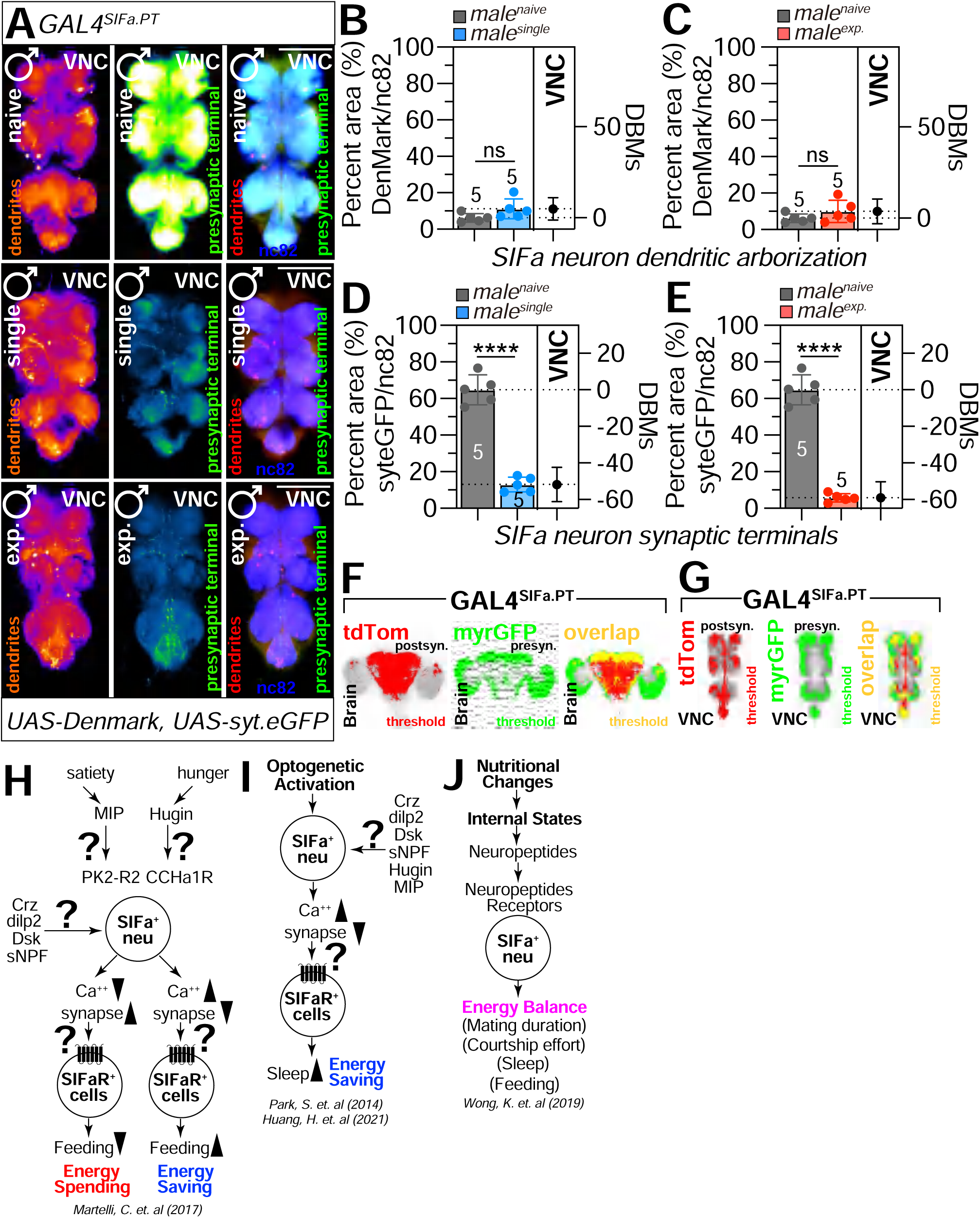
*SIFa* neurons project to important regions for control of reproductive behavior. (A) Distribution of dendrites and pre-synaptic terminals of neurons labeled via *GAL4^SIFa.PT^* in VNC in naïve (top panels), single (middle panels) and mated male flies (bottom panels). Flies expressing *GAL4^SIFa.PT^* together with *UAS-Denmark, UAS-syt.eGFP* were immunostained with anti-GFP (green), anti-DsRed (red) and nc82 (blue) antibodies. Scale bars represent 100 µm. (B-E) Quantification of dendritic arborization and synaptic terminals of *SIFa* neurons in Fig. A. (F-G) The threshold of RFP fluorescence (left panel, red), GFP fluorescence (middle panel, green) and overlapping area of GFP and RFP (right panel, yellow) in male fly brain and VNC was marked by threshold function of ImageJ. (H-J) Illustration depicting the association of *SIFa* neurons with the regulation of energy conservation and expenditure.

**Fig. S8.**
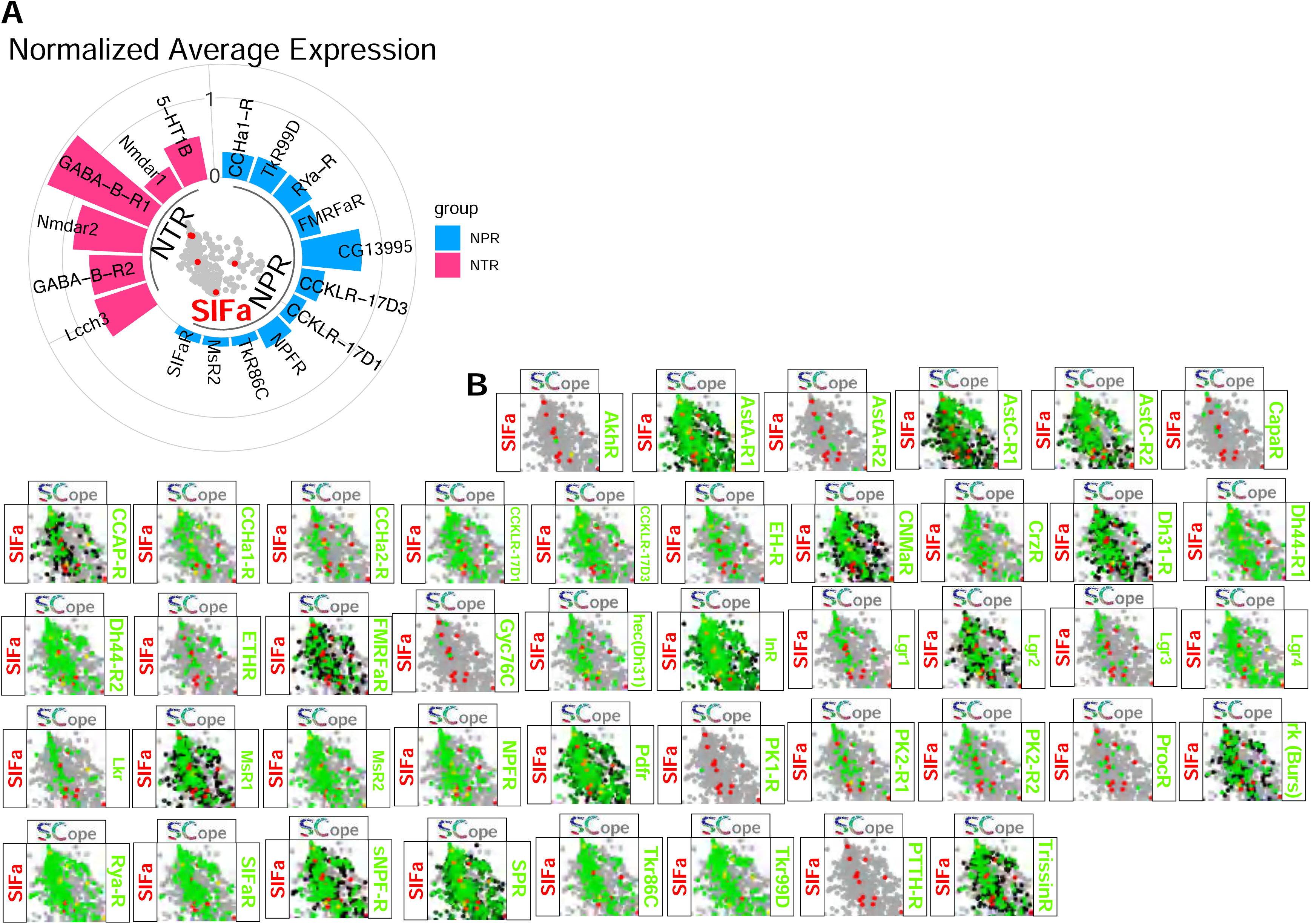
Bar plot of NPR and NTR expression in *SIFa+* cells. (A) The average UMI count was calculated for each gene in this population, followed by log^10^ transformation for data normalization. To focus on cells expressing these genes, only genes exhibiting increased expression of NTR and NPR were included in the bar plot. See the MATERIALS AND METHODS for a detailed description of the gene expression analysis used in this study. (B) Fly SCope single-cell RNA sequencing data of cells co-expressing *SIFa* together with neurotransmitter receptors and neuropeptide receptors.

## Supplementary information

**Document S1. Fig. S1–S5.** MD assay summary of NPs and NP-Rs-RNAi screening with *elav^c^*^155^ driver, related to Fig. 1.

(A-H’) MD assays for GAL4 mediated knockdown of AKH, AkhR, AstA, AstA-R2, MIP, AstC, AstC-R1, AstC-R2 (BL25940) (in the order of A-H) using the *elav^c^*^155^ driver.

(I-P’) MD assays for GAL4 mediated knockdown of AstC-R2 (BL36888), amn, CCHa1, CCHa1, CrZ, CrzR, CCAP-R, DH31-R (in the order of I-P) using the *elav^c^*^155^ driver.

(Q-X’) MD assays for GAL4 mediated knockdown of Dh44-R2, ETH-RNAi, ETHR, FMRFaR, ITP, ilp1, ilp2, ilp3 (in the order of Q-X) using the *elav^c^*^155^ driver.

(Y-f’) MD assays for GAL4 mediated knockdown of ilp4, ilp5, ilp6, ilp7, lp7, InR(31037), InR(31594), MsR2 (in the order of Y-f) using the *elav^c^*^155^ driver.

(g-n’) MD assays for GAL4 mediated knockdown of Nplp2, Proc-R, RYa-R, CCKLR-17D1, TkR86, 5-HT1A, 5-HT2A, 5-HT7 (in the order of g-n) using the *elav^c^*^155^ driver.

