## Supplementary figures and images for "Peptidergic neurons with extensive branching orchestrate the internal states and energy balance of male *Drosophila melanogaster*"

### 2024-SIFa-Document S1. Figures S1.pdf

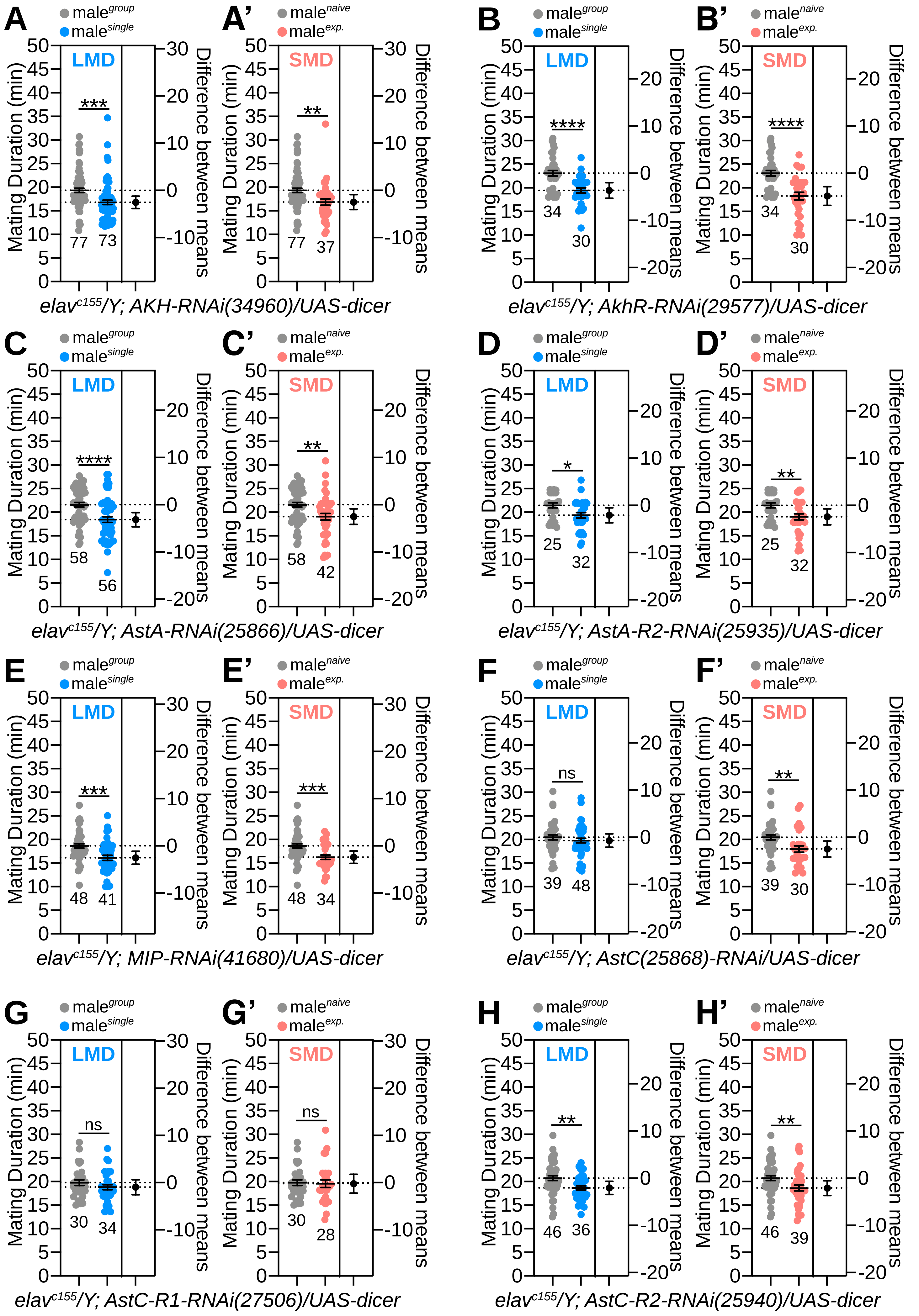

### 2024-SIFa-Document S1. Figures S2.pdf

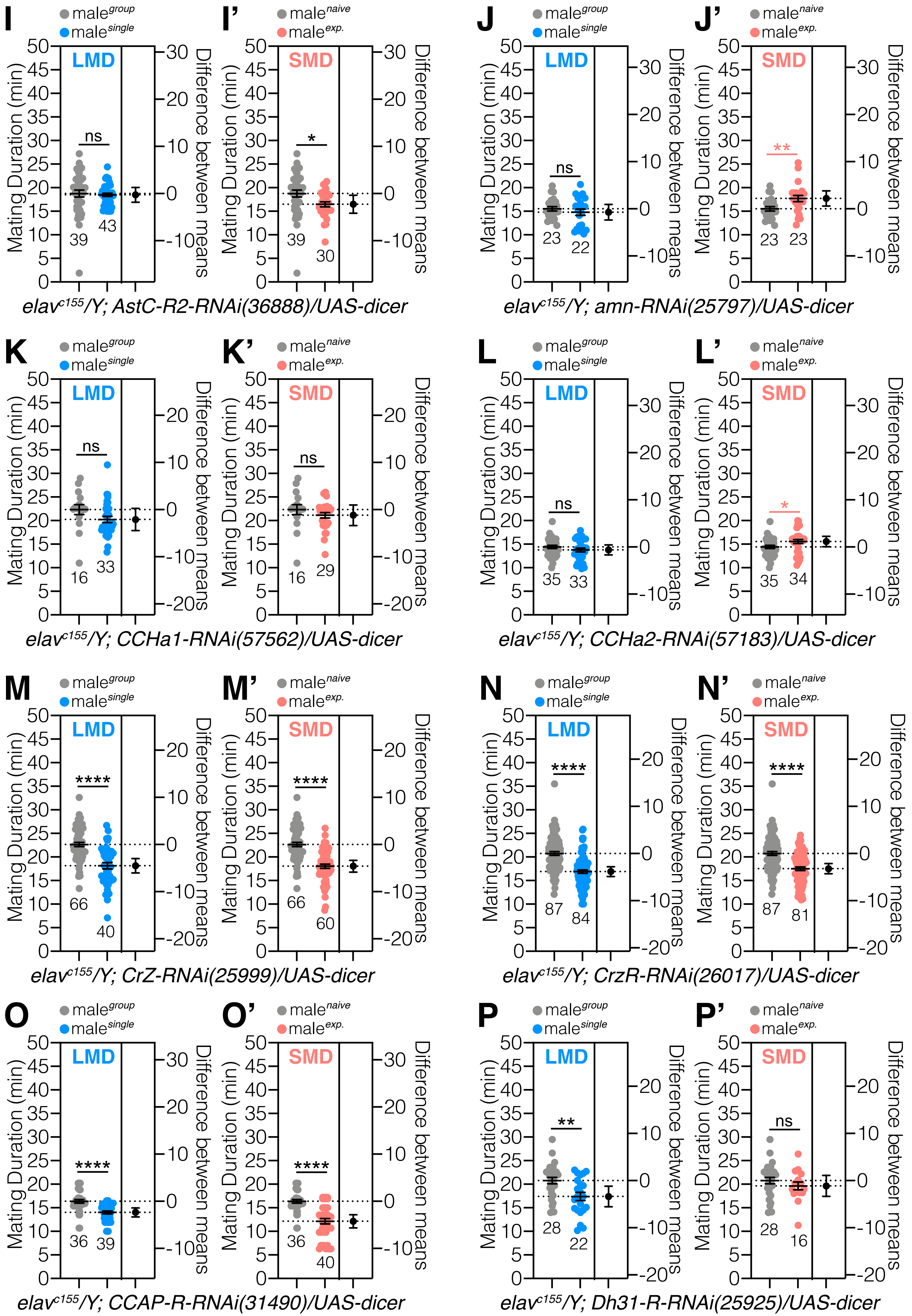

### 2024-SIFa-Document S1. Figures S3.pdf

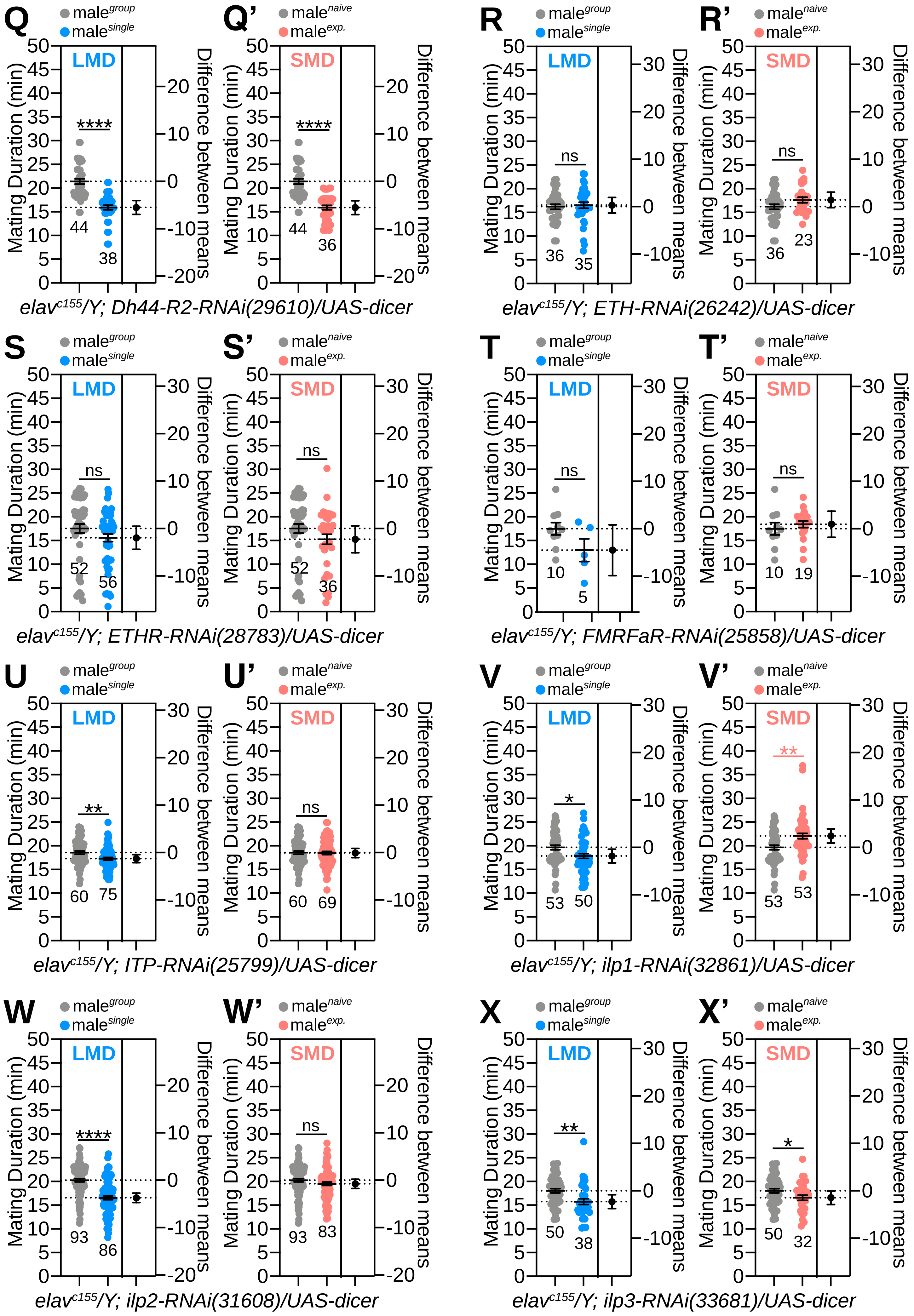

### 2024-SIFa-Document S1. Figures S4.pdf

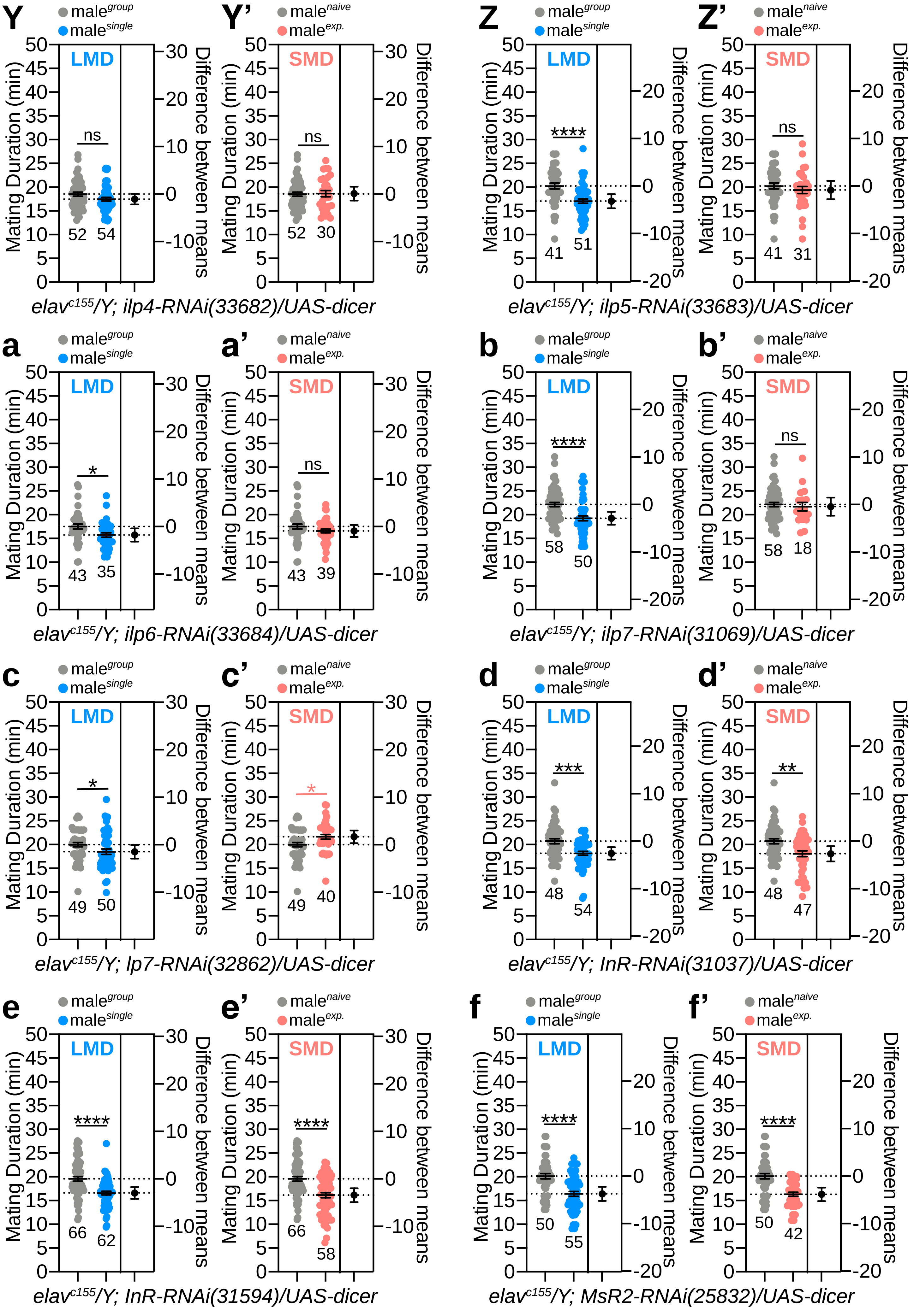

### 2024-SIFa-Document S1. Figures S5.pdf

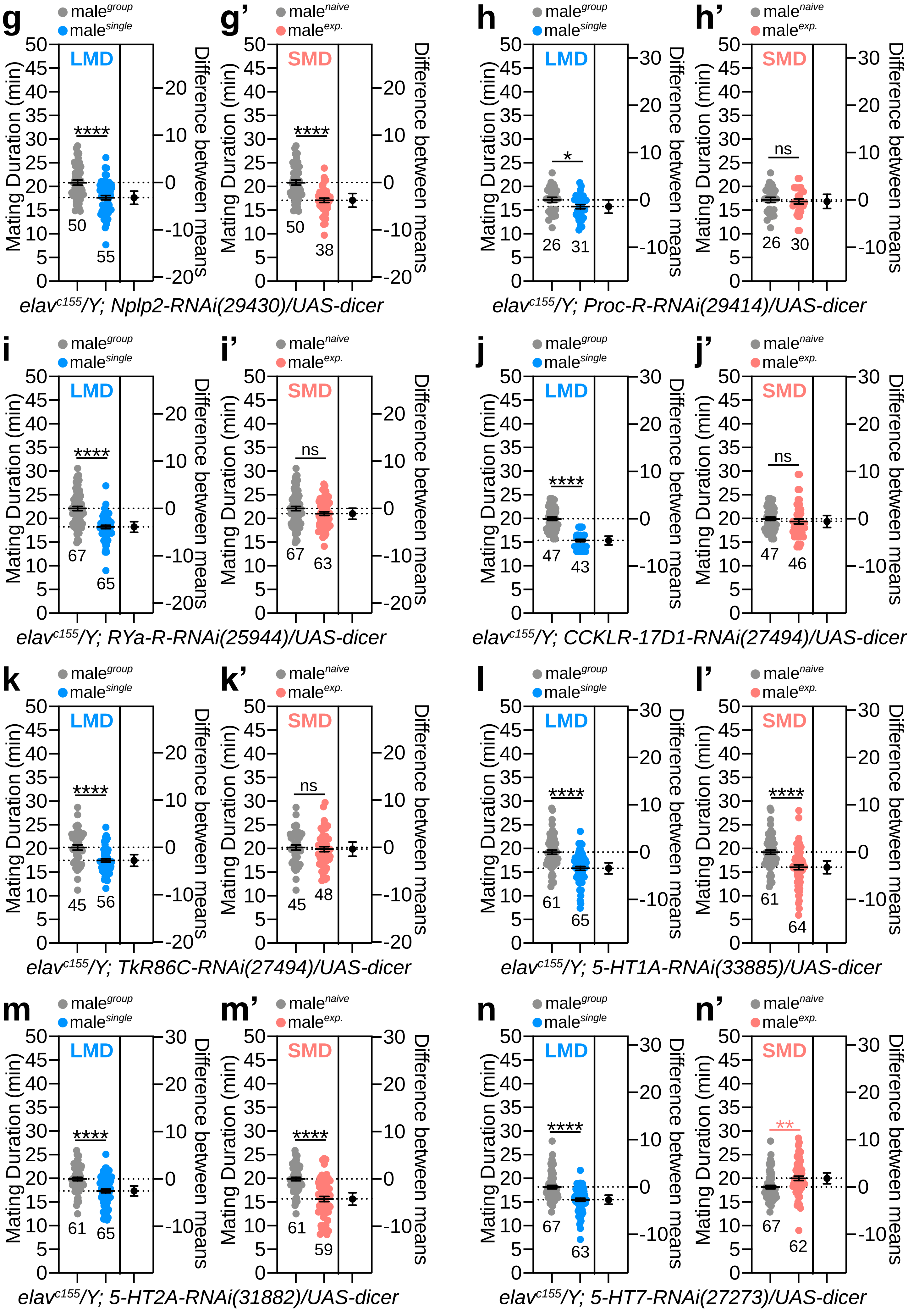
