## Supplement Table 1 for "Peptidergic neurons with extensive branching orchestrate the internal states and energy balance of male *Drosophila melanogaster*": Song_2024_SIFa_Table.S1.pdf

**Table S1.** Summary of neuropeptide receptors *RNAi* screening with *elav*$^{c155}$ driver

| RNAi /stock # | naive | single | exp. | LMD | | | SMD | | |
|---|---|---|---|---|---|---|---|---|---|
| | *n* | *n* | *n* | *t* | *df* | *p-value* | *t* | *df* | *p-value* |
| *AKH-RNAi* 34960 | 77 | 73 | 37 | 3.827 | 148 | 0.0002 | 3.084 | 112 | 0.0026 |
| *AkhR-RNAi* 29577 | 34 | 30 | 30 | 4.419 | 62 | <0.0001 | 4.881 | 62 | <0.0001 |
| *AstA-RNAi* 25866 | 58 | 56 | 42 | 4.145 | 112 | <0.0001 | 3.054 | 98 | 0.0029 |
| *AstA-R2-RNAi* 25935 | 25 | 32 | 32 | 2.651 | 55 | 0.0105 | 2.855 | 55 | 0.0061 |
| *MIP-RNAi* 49087 | 48 | 41 | 34 | 3.651 | 87 | 0.0004 | 3.620 | 80 | 0.0005 |
| *AstC-RNAi* 51988 | 39 | 48 | 30 | 0.9749 | 85 | 0.3324 | 2.925 | 67 | 0.0047 |
| *AstC-R1-RNAi* 27506 | 30 | 34 | 28 | 1.193 | 62 | 0.2373 | 0.2198 | 56 | 0.8268 |
| *AstC-R2-RNAi* 25940 | 46 | 36 | 39 | 2.954 | 80 | 0.0041 | 2.657 | 83 | 0.0095 |
| *AstC-R2-RNAi* 36888 | 39 | 43 | 30 | 0.3416 | 80 | 0.7335 | 2.377 | 67 | 0.0203 |
| *amn-RNAi* 25797 | 23 | 22 | 23 | 0.8960 | 43 | 0.3752 | 2.757 | 44 | 0.0085* |
| *CCHa1-RNAi* 57652 | 16 | 33 | 29 | 1.784 | 47 | 0.0809 | 1.100 | 43 | 0.2776 |
| *CCHa2-RNAi* 57183 | 35 | 33 | 34 | 1.198 | 66 | 0.2352 | 2.104 | 67 | 0.0391* |
| *CRZ-RNAi* 25999 | 66 | 40 | 60 | 5.817 | 104 | <0.0001 | 7.288 | 124 | <0.0001 |
| *CrzR-RNAi* 26017 | 87 | 84 | 81 | 7.192 | 169 | <0.0001 | 5.892 | 166 | <0.0001 |
| *CCAP-R-RNAi* 31490 | 36 | 39 | 40 | 4.896 | 73 | <0.0001 | 6.029 | 74 | <0.0001 |
| *Dh31-R-RNAi* 25925 | 28 | 22 | 16 | 3.120 | 48 | 0.0031 | 1.027 | 42 | 0.3104 |
| *Dh44-R2-RNAi* 51974 | 44 | 38 | 36 | 7.441 | 80 | <0.0001 | 7.513 | 78 | <0.0001 |
| *ETH-RNAi* 26242 | 36 | 35 | 23 | 0.3755 | 69 | 0.7085 | 1.732 | 57 | 0.0886 |
| *ETHR-RNAi* 28783 | 52 | 56 | 36 | 1.599 | 106 | 0.1128 | 1.592 | 86 | 0.1150 |
| *FMRFaR-RNAi* 25858 | 10 | 5 | 19 | 1.820 | 13 | 0.0919 | 0.7027 | 27 | 0.4882 |
| *ITP-RNAi* 25799 | 60 | 75 | 69 | 3.010 | 133 | 0.0031 | 0.1614 | 127 | 0.8721 |
| *ilp1RNAi* 32861 | 53 | 50 | 53 | 2.524 | 101 | 0.0132 | 3.237 | 104 | 0.0016* |
| *ilp2-RNAi* 31608 | 93 | 86 | 83 | 7.348 | 177 | <0.0001 | 1.546 | 174 | 0.1240 |

*- Crosses showed effect on LMD/SMD behaviors are colored as sky blue (LMD defect) or pink (SMD defect).*
** The mating duration of each condition reverses due to defective LMD or SMD.*

*Table 1 Continued*

| RNAi /stock # | naive | single | exp. | LMD | | | SMD | | |
|---|---|---|---|---|---|---|---|---|---|
| | *n* | *n* | *n* | *t* | *df* | *p-value* | *t* | *df* | *p-value* |
| *ilp3-RNAi 33681* | 50 | 38 | 32 | 3.136 | 86 | 0.0023 | 2.058 | 80 | 0.0429 |
| *ilp4-RNAi 33682* | 52 | 54 | 30 | 1.934 | 104 | 0.0558 | 0.1304 | 80 | 0.8965 |
| *ilp5-RNAi 33683* | 41 | 51 | 31 | 4.279 | 90 | <0.0001 | 0.8735 | 70 | 0.3854 |
| *ilp6-RNAi 33684* | 43 | 35 | 39 | 2.562 | 76 | 0.0124 | 1.467 | 80 | 0.1463 |
| *ilp7-RNAi 31069* | 58 | 50 | 18 | 4.279 | 106 | <0.0001 | 0.4707 | 74 | 0.6392 |
| *ilp7-RNAi 32862* | 49 | 50 | 40 | 2.042 | 97 | 0.0439 | 2.480 | 87 | 0.0151* |
| *InR-RNAi 31037* | 48 | 54 | 47 | 3.855 | 100 | 0.0002 | 3.262 | 93 | 0.0015 |
| *InR-RNAi 31594* | 66 | 62 | 58 | 4.753 | 126 | <0.0001 | 4.691 | 122 | <0.0001 |
| *MsR2-RNAi 25832* | 50 | 55 | 42 | 5.015 | 103 | <0.0001 | 5.397 | 90 | <0.0001 |
| *Nplp2-RNAi 29430* | 50 | 55 | 38 | 4.544 | 103 | <0.0001 | 5.184 | 86 | <0.0001 |
| *Proc-R-RNAi 29414* | 26 | 31 | 30 | 2.010 | 55 | 0.0494 | 0.4073 | 54 | 0.6854 |
| *Rya-R-RNAi 25944* | 67 | 65 | 63 | 6.933 | 130 | <0.0001 | 1.887 | 128 | 0.0614 |
| *CCKLR-17D1 25999* | 47 | 43 | 46 | 9.877 | 88 | <0.0001 | 0.8616 | 91 | 0.3911 |
| *TkR86C-RNAi 27494* | 45 | 56 | 48 | 4.417 | 99 | <0.0001 | 0.4682 | 91 | 0.6408 |
| *5-HT1A-RNAi 33885* | 61 | 65 | 64 | 5.781 | 124 | <0.0001 | 4.677 | 123 | <0.0001 |
| *5-HT2A-RNAi 25925* | 61 | 66 | 59 | 4.768 | 125 | <0.0001 | 6.434 | 118 | <0.0001 |
| *5-HT7--RNAi 27273* | 67 | 63 | 62 | 5.552 | 128 | <0.0001 | 3.208 | 127 | 0.0017* |

*- Crosses showed effect on LMD/SMD behaviors are colored as sky blue (LMD defect) or pink (SMD defect).*
** The mating duration of each condition reverses due to defective LMD or SMD.*
